# Mitochondrial-targeted plastoquinone therapy ameliorates early onset muscle weakness that precedes ovarian cancer cachexia in mice

**DOI:** 10.1101/2024.10.22.619751

**Authors:** Luca J. Delfinis, Shahrzad Khajehzadehshoushtar, Luke D. Flewwelling, Nathaniel J. Andrews, Madison C. Garibotti, Shivam Gandhi, Aditya N. Brahmbhatt, Brooke A. Morris, Bianca Garlisi, Sylvia Lauks, Caroline Aitken, Stavroula Tsitkanou, Jeremy A. Simpson, Nicholas P. Greene, Arthur J. Cheng, Jim Petrik, Christopher G.R. Perry

## Abstract

Cancer cachexia, and the related loss of muscle and strength, worsens quality of life and lowers overall survival. Recently, a novel ‘pre-atrophy’ muscle weakness was identified during early-stage cancer. While mitochondrial stress responses are associated with early-stage pre-atrophy weakness, a causal relationship has not been established. Using a robust mouse model of metastatic epithelial ovarian cancer (EOC)-induced cachexia, we found the well-established mitochondrial-targeted plastoquinone SkQ1 partially prevents pre-atrophy weakness in the diaphragm. Furthermore, SkQ1 improved force production during atrophy without preventing atrophy itself in the tibialis anterior and diaphragm. EOC reduced flexor digitorum brevis (FDB) force production and myoplasmic free calcium ([Ca^2+^]_i_) during contraction in single muscle fibers, both of which were prevented by SkQ1. Remarkably, changes in mitochondrial reactive oxygen species and pyruvate metabolism were heterogeneous across time and between muscle types which highlights a considerable complexity in the relationships between mitochondria and muscle remodeling throughout EOC. These discoveries identify that muscle weakness can occur independent of atrophy throughout EOC in a manner that is linked to improved calcium handling. The findings also demonstrate that mitochondrial-targeted therapies exert a robust effect in preserving muscle force during the early pre-atrophy period and in late-stage EOC once cachexia has become severe.

## Introduction

Cancer-induced cachexia is a multifactorial syndrome characterized by muscle atrophy and functional impairment (1). This condition leads to reductions in quality of life, tolerance to anticancer therapies and overall survivability (2–4). Although dozens of clinical investigations have been completed for cancer cachexia therapy development, none have produced sufficiently positive results to yield a therapeutic intervention (5). This could be due to a mismatch between clinical reality and current animal models, resulting in poor translation of preclinical studies (6). Indeed, most animal models have been considered insufficient due to the lack of metastasis or ectopic tumour growth (tumour growth that does not occur in host organ), among other reasons. Therefore, more translationally powerful orthotopic, and metastatic models are believed to improve the predictive power of preclinical models in terms of both mechanism and therapy elucidation for cancer cachexia (6).

While most research has focused on the mechanisms of muscle atrophy in cancer cachexia, our group recently reported muscle weakness precedes atrophy and whole-body cachexia (7, 8). We showed this in both limb and respiratory muscles within the C26 model of colorectal cancer (7) and a novel model of epithelial ovarian cancer (EOC)_that demonstrates the clinically relevant characteristic of metastasis (8). These discoveries suggest pre-atrophy weakness could be a ubiquitous phenomenon across muscles and cancer types. Moreover, while previous studies show that mitochondrial dysregulation precedes atrophy in cancer cachexia (9), we demonstrated mitochondrial stress coincides with pre-atrophy weakness in two models of cancer cachexia (7, 8). More specifically, we demonstrated early weakness was related to lower pyruvate oxidation and attenuated gene programs related to oxidative phosphorylation, as well as elevated reactive oxygen species (ROS). These alterations also depended on muscle type (8). Other groups have shown the mitochondrial-targeted compounds MitoQ (containing ubiquinone) and SS-31 (mitochondrial cardiolipin-targeting small peptide) attenuate atrophy in cell culture and animal models to some effect and, in the case of SS-31, preserve muscle force generating capacity in relation to certain indices of mitochondrial function in the C26 model (10, 11). However, the degree to which mitochondrial stress responses contribute to pre-atrophy weakness vs atrophy itself remains unresolved. Likewise, it remains unknown if mitochondria exerts stressors that contribute to weakness once atrophy occurs but through mechanisms that are not driven by atrophy itself.

Considering ovarian cancer-induced cachexia is relatively understudied and is the most lethal gynecological cancer in women (12), we aimed to explore this topic in our recently reported novel orthotopic model of metastatic ovarian cancer cachexia that is inducible in adult mice (8). The purpose of this investigation was to determine the degree to which pharmacological targeting of mitochondria can prevent i) the recently identified pre-atrophy weakness that occurs during cancer, ii) weakness during atrophy, and iii) atrophy itself. We employed a lipophilic (tissue penetrating) cationic (mitochondrial-targeting) plastoquinone-based antioxidant, SkQ1, that is known to prevent electron slip from the electron transport chain (13). We found muscle force production is partially preserved prior to the onset of atrophy as well as once atrophy has occurred without preserving muscle fiber cross sectional area. These findings demonstrate that mitochondria are contributors to weakness that occurs independent of a loss of muscle mass during ovarian cancer.

## Results

### SkQ1 administration in drinking water does not affect mouse survival, body weight, fat weight, tumour weight, muscle wet weight or 4-HNE protein adducts

C57BL6J mice were injected at 10 and 16 weeks of age with 1 million EOC cells and treated with SkQ1 or water to develop Early-Stage (Early-EOC CON and Early-EOC SkQ1) and Late-Stage (Late-EOC CON and Late-EOC SkQ1) cancer timepoints **(Figure 1A)**. The Late-Stage timepoint was characterized by severe metastasis on the abdominal surface of the diaphragm and extensive ascites (58-82 days post EOC injection; **Figure 1B and SFigure 1A)**, while the Early-Stage was characterized by no apparent metastasis or ascites (35-46 days post EOC injection**; SFigure 1A).** There were no statistical differences in Kaplan-Meier estimates of survival between Late-EOC SkQ1 and Late-EOC CON treated mice **(Figure 1C)** and no statistical difference in the days alive post-EOC inoculation, suggesting SkQ1 did not improve survival **(Figure 1D)**. There were no differences in body weight over time between the Sham group and Early-Stage mice as well as Sham and Late-Stage mice regardless of SkQ1 or water treatments **(Figure 1E).** Moreover, there were no differences in final body weight of Sham vs Early-Stage mice, but there was a significantly lower final body weight in both SkQ1 and CON Late-Stage mice compared to Sham indicative of cachexia development in Late-Stage cancer **(Figure 1F).** Recognizing that cachexia lowers body weight, we also compared the change in weight from peak values (regardless of when they occurred) to final values. There was a significant decrease in percent body weight change from peak to final body weight in both Early-Stage (-8% in Early-EOC CON and -5% in Early-EOC SkQ1) and Late-Stage mice (-33% in Late-EOC CON and Late-EOC SkQ1) compared to Sham **(Figure 1G)**. Mice at the Early-Stage time point exhibited no changes in subcutaneous adipose mass in the inguinal fat pad compared to Sham, but there were significant reductions in Late-Stage mice compared to Sham **(Figure 1H).** There were also no statistical differences in estimated average daily food consumption between any groups in Early-Stage or Late-Stage mice **(SFigure 1B)**, albeit food consumption calculations per animal were estimated as mice were group housed and food weights were taken once a week. Similarly, there were no differences in estimated daily water consumption in Sham vs Early- or Late-Stage mice **(SFigure 1C).** There was a significant increase in primary ovarian tumour mass at both the Early-Stage and Late-Stage timepoints compared to Sham **(Figure 1I)**. This finding also suggests SkQ1 did not affect tumour size. Also, spleen mass (marker of inflammation) was not changed at the Early- and Late-Stage timepoints compared to Sham although Late-Stage groups demonstrated more variability compared to other groups **(SFigure 1D).** There were no reductions in hindlimb muscle wet weights in Early-Stage mice compared to Sham, but significant reductions in soleus (SOL), extensor digitorum longus (EDL), plantaris (PLT), TA, gastrocnemius (GA) and quadriceps (QUAD) skeletal muscle in Late-Stage mice compared to Sham **(Figure 1J)**. We also evaluated volitional wheel running distance after 24 hours of exposure to a running wheel and found no differences in Sham vs Early-Stage mice **(SFigure 1E)**. We were unable to assess volitional wheel running in Late-Stage mice given the challenges in predicting mouse survival 48 hours prior to tissue collection in this advanced cancer time point.

**Figure 1.**
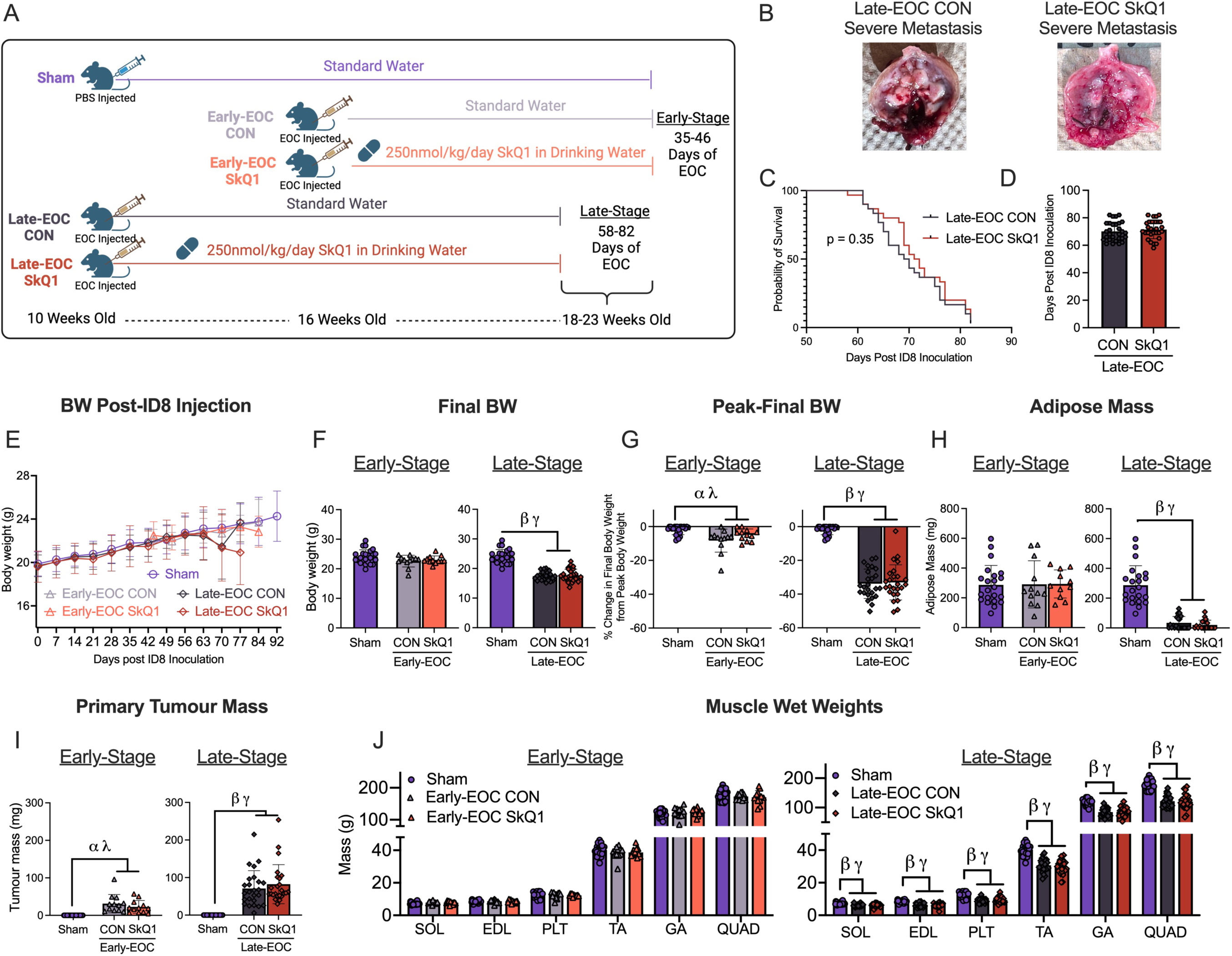
SkQ1 treatment in drinking water does not change survivability, body weight loss, adipose mass, tumour mass or muscle wet weights at Early- or Late-stage tumour development in a metastatic ovarian cancer cachexia mouse model. 1 x 10^6^ ID8 cells were injected underneath the ovarian bursa and developed for 35-46 or 58-82 days (Early-Stage and Late-Stage respectively). Mice were treated with standard drinking water (Early-EOC CON and Late-EOC CON) or 250nmol/kg/day SkQ1 (Early-EOC SkQ1 and Late-EOC SkQ1). Control mice were sham injected with identical volumes of PBS and developed for 91-105 days (Sham) **(A).** Metastasis is imaged in Late-Stage mice **(B).** Kaplan-Meier probability of survivability and days alive post cancer injection analysis (**C, D** n=30). Body weight over time (**E**, n=12-27) Final body weight (**F**, n=12-27) and percent change from peak to final body weight (**G**, n=12-27) was assessed. Subcutaneous adipose mass from the inguinal fat pad (**H**, n=11-23), primary ovarian tumours (**I**, n=12-25) and muscle wet weights (**J**, n=12-25) were also analyzed. Figure C was analyzed with a Simple Survival Analysis (Kaplan-Meier), Figure D with an unpaired T-Test. All other data was analyzed using a one-way ANOVA and followed by a two-stage step-up method of Benjamini, Krieger and Yukutieli multiple comparisons test. Data that was not normally distributed was analyzed with a Kruskal-Wallis test followed by the same post-hoc analysis. Sham Control (Sham), Early-Stage Cancer Control (Early-EOC CON), Early-Stage Cancer SkQ1 (Early-EOC SkQ1), Late-Stage Cancer Control (Late-EOC CON), Late-Stage Cancer SkQ1 (Late-EOC SkQ1), Body weight (BW), soleus (SOL), extensor digitorum longus (EDL), plantaris (PLT), tibialis anterior (TA), gastrocnemius (GA), quadriceps (QUAD). Results represent mean ± SD. *α P*<0.05 Sham Vs Early-EOC CON; *λ P*<0.05 Sham Vs Early-EOC SkQ1; *β P*<0.05 Sham Vs Late-EOC CON; *γ P*<0.05 Sham Vs Late-EOC SkQ1.

### SkQ1 administration in drinking water lowers mRNA content of Atrogin, interleukin -6 (IL6) and tumour necrosis factor (TNF)

We evaluated the mRNA content of canonical markers of muscle atrophy and inflammation in the TA and diaphragm muscles at both cancer time points (14). These muscles were selected given we recently reported their time-dependent responses to cancer in this model. In the TA, there were no changes in mRNA content of *Il6* or *Tnf* at both timepoints **(SFigure2A and 2B)**. Interestingly, SkQ1 reduced cancer-induced increases in mRNA content of *Atrogin* in Late-Stage cancer mice **(*P* = 0.05, SFigure 2C)**, but did not reduce cancer-induced elevated levels of *Murf1* **(SFigure 2D)**. In the diaphragm, SkQ1 reduced cancer-induced increases in *Il6* mRNA content in Early-Stage and Late-Stage mice **(*P* = 0.08, SFigure 2E)**. mRNA content of *Tnf* was not different at Early-Stage but SkQ1 reduced Late-Stage cancer-induced increases in *Tnf* mRNA content **(*P* = 0.07, SFigure 2F)**. *Atrogin* mRNA contents were unchanged at the Early-Stage time point but was interestingly decreased in Late-EOC CON and Late-EOC SkQ1 mice **(SFigure 2G)**. Last, *Murf1* mRNA content was unchanged at the Early-Stage and Late-Stage time points in the diaphragm **(SFigure 2H)**.

### SkQ1 administration in drinking water attenuates cancer-induced muscle weakness in the TA and diaphragm at Early-Stage in addition to diaphragm weakness in Late-Sage without attenuating atrophy

TA force production *in-situ* was decreased in Early-EOC CON compared to Sham (**Figure 2A**). However, SkQ1 prevented 46% of this weakness **(Figure 2A and 2B)**. This indicates that SkQ1 treatment improved force production in the TA during the Early-Stage timepoint. There were no differences in rate of contraction and time to relaxation at 100Hz stimulation frequency in Early-Stage mice **(Figure 2C and 2D).** Similar conclusions were made regarding absolute force in the TA at early-stage (data not shown). We were unable to collect force production at the Late-Stage time point as we were not able to perform *in-situ* force analyses due to their advanced frailty. Early-EOC CON showed lower CSA in MHCIIa whereas both Early-EOC CON and Early-EOC SkQ1 showed lower CSA in MHCIIx and pooled fibers compared to Sham **(Figure 2E and 2F)**. There was no statistical difference in MHCIIa CSA between Sham and Early-EOC SkQ1. This suggests that SkQ1 treatment improved force production without affecting fiber CSA in the Early-Stage timepoint. Last, Late-EOC CON and Late-EOC SkQ1 mice exhibited decreases in all MHC isoforms (MHCIIa, MHCIIx and MHCIIb) compared to Sham but there was no difference between CON and SkQ1 at this time point **(Figure 2G-I).**

**Figure 2.**
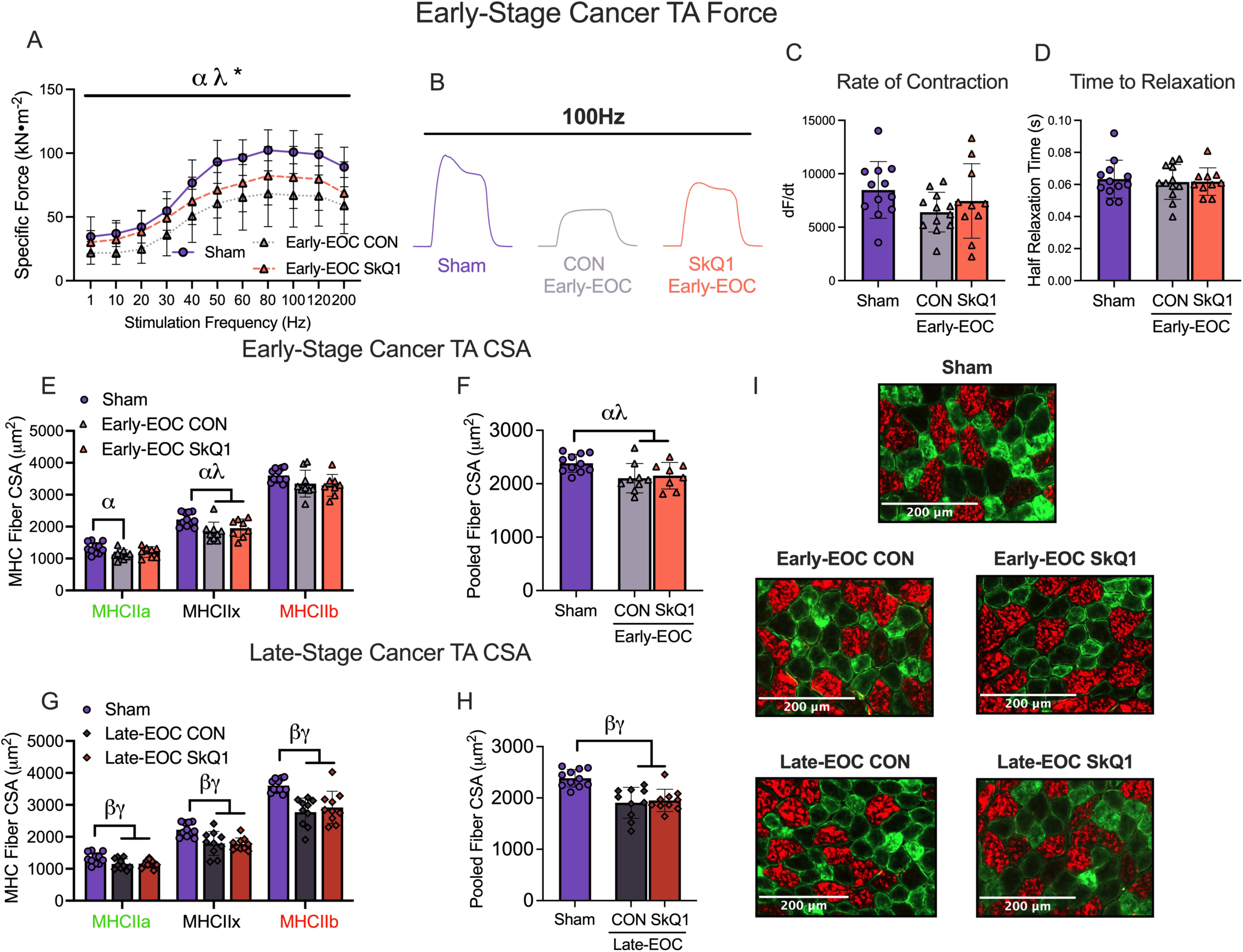
SkQ1 treatment in drinking water attenuates Early-Stage muscle weakness but does not change Early- or Late-Stage fiber cross-sectional area in tibialis anterior muscle in a metastatic ovarian cancer cachexia mouse model. In situ TA force production was assessed using the force-frequency relationship (**A** n=10-12; **B** representative twitches at 100Hz). Rate of twitch contraction along with the half relaxation time was also analyzed at 100Hz (**C, D** n=10-12). Analysis of fiber histology on MHC isoforms was performed in all groups. CSA of MHC isoforms were evaluated in the TA along with pooled fibers (**E-H**, n =8-11; **I** representative images at 20x magnification). A two-way ANOVA was used for figure A. All other data was analyzed using a one-way ANOVA and followed by a two-stage step-up method of Benjamini, Krieger and Yukutieli multiple comparisons test. Data that was not normally distributed was analyzed with a Kruskal-Wallis test followed by the same post-hoc analysis. Sham Control (Sham), Early-Stage Cancer Control (Early-EOC CON), Early-Stage Cancer SkQ1 (Early-EOC SkQ1), Late-Stage Cancer Control (Late-EOC CON), Late-Stage Cancer SkQ1 (Late-EOC SkQ1), CSA (cross-sectional area), MHC (myosin heavy chain), TA (tibialis anterior). Results represent mean ± SD. *α P*<0.05 Sham Vs Early-EOC CON; *λ P*<0.05 Sham Vs Early-EOC SkQ1; * *P*<0.05 Early-EOC CON Vs Early-EOC SkQ1 *β P*<0.05 Sham Vs Late-EOC CON; *γ P*<0.05 Sham Vs Late-EOC SkQ1.

Diaphragm force production assessed *in vitro* was decreased in Early-EOC CON and Early-EOC SkQ1 mice compared to Sham **(Figure 3A and 3B)**. However, Early-EOC SkQ1 mice exhibited an average 31% higher force production across all stimulation frequencies relative to Early-EOC CON mice **(Figure 3A and 3B)** preventing 62% of cancer-induced muscle weakness. This demonstrates that SkQ1 treatment also improves force production in the Early-Stage timepoint within the diaphragm. There were no differences in rate of contraction and time to relaxation at 100Hz stimulation frequency between Sham and Ealy-EOC CON or Early-EOC SkQ1 **(Figure 3C and 3D).** This same relationship existed at the Late-Stage timepoint such that Late-EOC CON mice exhibited decrease in muscle force production which was partially improved in Late-EOC SkQ1 mice (average improvement of 41% across all frequencies, preventing 30% of cancer-induced weakness) with no changes in rate of contraction or time to relaxation at 100Hz stimulation frequency **(Figure 3E – 3H)**. There was no effect of SkQ1 on the development of fatigue at Early-Stage or Late-Stage cancer development within the diaphragm **(SFigure 3A and 3B)**. Similar to the TA, SkQ1 treatment did not change fiber CSA in the diaphragm **(Figure 3I-3M)**.

**Figure 3.**
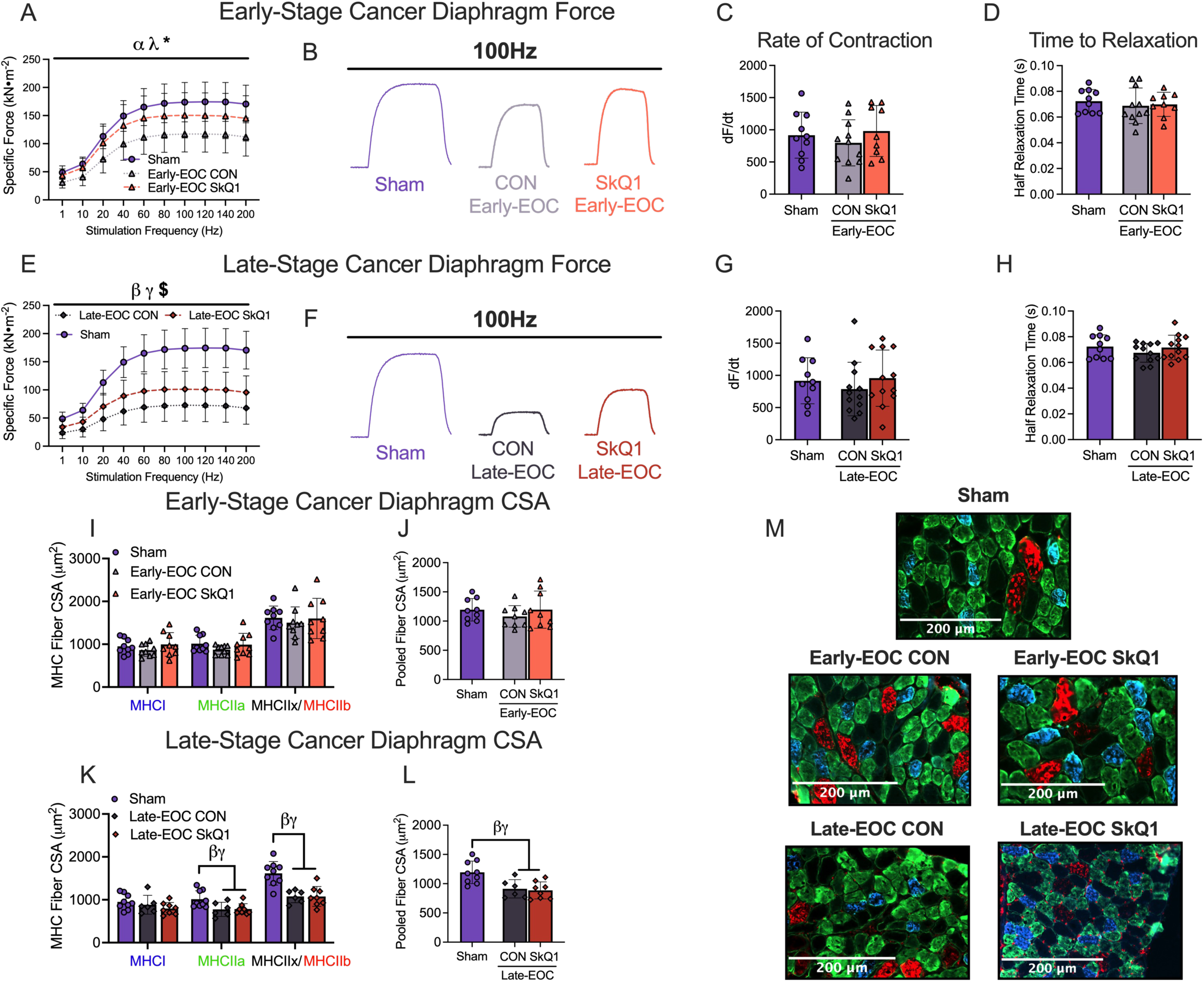
SkQ1 treatment in drinking water attenuates Early- and Late-stage muscle weakness but does not change Early- or Late-stage fiber cross-sectional area in diaphragm muscle in a metastatic ovarian cancer cachexia mouse model. In Vitro diaphragm force production was assessed using the force-frequency relationship (**A & E** n=9-12; **B & F** representative twitches at 100Hz). Rate of twitch contraction along with the half relaxation time was also analyzed at 100Hz (**C, D & G, H** n=10-12). Analysis of fiber histology on MHC isoforms was performed in all groups. CSA of MHC isoforms were evaluated in the diaphragm along with pooled fibers (**I-L**, n =6-9; **I** representative images at 20x magnification). A two-way ANOVA was used for figures A and E. All other data was analyzed using a one-way ANOVA and followed by a two-stage step-up method of Benjamini, Krieger and Yukutieli multiple comparisons test. Data that was not normally distributed was analyzed with a Kruskal-Wallis test followed by the same post-hoc analysis. Sham Control (Sham), Early-Stage Cancer Control (Early-EOC CON), Early-Stage Cancer SkQ1 (Early-EOC SkQ1), Late-Stage Cancer Control (Late-EOC CON), Late-Stage Cancer SkQ1 (Late-EOC SkQ1), CSA (cross-sectional area), MHC (myosin heavy chain). Results represent mean ± SD. *α P*<0.05 Sham Vs Early-EOC CON; *λ P*<0.05 Sham Vs Earl-EOC SkQ1; * *P*<0.05 Early-EOC CON Vs Early-EOC SkQ1; *β P*<0.05 Sham Vs Late-EOC CON; *γ P*<0.05 Sham Vs Late-EOC SkQ1; $ *P*<0.05 Late-EOC CON Vs Late-EOC SkQ1.

Collectively, in the diaphragm, Early-Stage mice showed muscle weakness without atrophy, which is improved by SkQ1 treatment. In Late-Stage mice, where muscle weakness and atrophy coexisted, SkQ1 treatment enhanced force production independently of CSA. These results further support that muscle weakness can be caused by mechanisms distinct from atrophy.

### SkQ1 administration in drinking water lowers Complex I-supported mH_2_O_2_ but also decreases Complex I-supported mitochondrial respiration in the TA

We also evaluated mitochondrial respiration and mH_2_O_2_ considering SkQ1 prevents electron slip in certain components of the electron transport chain (13). Using permeabilized muscle fibers, we stimulated complex I-supported mH_2_O_2_ with forward electron transfer (pyruvate and malate) to generate NADH (15) **(Figure 4A).** Substrate-specific maximal mH_2_O_2_ kinetics (State II) were followed by titration of ADP to determine the ability of ADP to attenuate mH_2_O_2_ during oxidative phosphorylation (State III). In the TA at the Early-Stage timepoint, there were no differences in maximal mH_2_O_2_ nor after the addition of ADP in absolute terms (reflective of rates during oxidative phosphorylation) or after expressing the effect of ADP relative to maximal mH_2_O_2_ (indicative of mitochondrial sensitivity to ADP) **(Figure 4B-D)**. At the Late-Stage timepoint, there was a significant decrease in maximal mH_2_O_2_ in Late-EOC CON mice compared to Sham but not in Late-EOC SkQ1 mice compared to Sham **(Figure 4E).** Despite the lower maximal mH_2_O_2_ in Late-EOC CON, which is an artificial condition in the absence of ADP-stimulated ATP synthesis, there was a significant increase in mH_2_O_2_ in Late-EOC CON and Late-EOC SkQ1 mice compared to Sham after titration of ADP, suggestive of increased absolute rates of mH_2_O_2_ across a wide range of oxidative phosphorylation kinetics **(Figure 4F)**. To determine if these changes in absolute rates in the presence of ADP were due to changes in mitochondrial responsiveness to ADP, we then expressed mH_2_O_2_ at each [ADP] relative to maximal mH_2_O_2_ as a measure of the relative effect of ADP. This analyses revealed that that mH_2_O_2_ at a given [ADP] relative to maximal mH_2_O_2_ is also elevated in Late-EOC CON similar to the pattern in absolute rates **(Figure 4G)**. Collectively, these findings demonstrate that reduced mitochondrial ADP sensitivity is a specific mechanism by which cancer elevates TA mitochondrial mH_2_O_2_ in Late-EOC CON. This relative analysis also revealed that Late-EOC SkQ1 mice exhibited decreases in mH_2_O_2_ compared to Late-EOC CON mice **(Figure 4G)**. This suggests that SkQ1 improves the ability of ADP to suppress mH_2_O_2_ as a specific mechanism underlying its antioxidant effect in this model, at least in regard to complex I-supported mH_2_O_2_ during oxidative phosphorylation. We also measured 4HNE as a marker of global membrane redox modification to explore a potential mechanism of decreased oxidative stress with SkQ1 treatment. We found no changes in the content of 4HNE adducts in the TA in Early-Stage mice, but unexpectedly found a decrease in Late-Stage EOC CON and SkQ1 mice vs Sham **(SFigure 5A)**.

**Figure 4.**
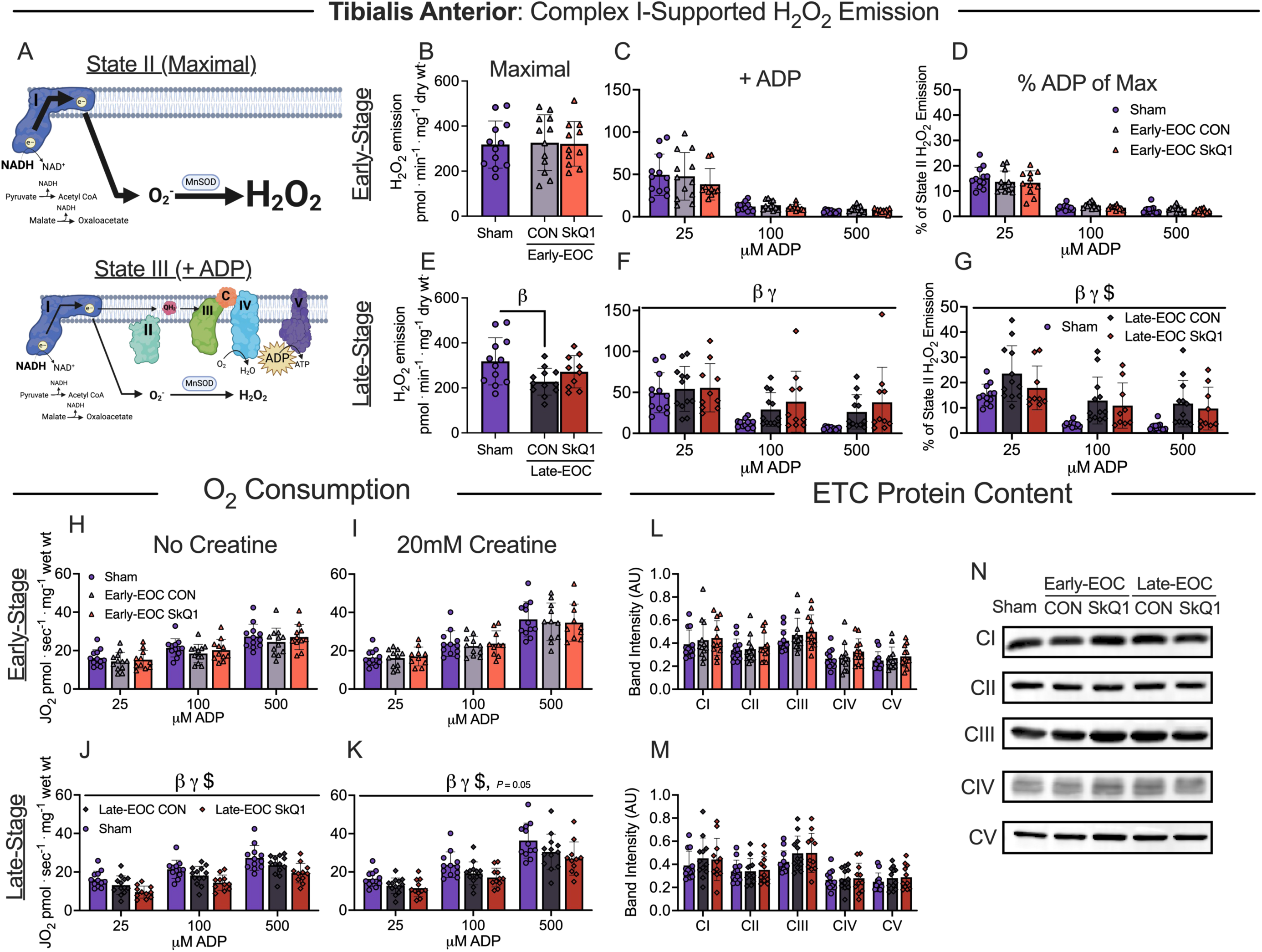
SkQ1 treatment in drinking water reduces Late-Stage Complex I stimulated H_2_O_2_ and, ADP-supported mitochondrial respiration in tibialis anterior muscle within a metastatic ovarian cancer cachexia mouse model. Complex I stimulated H_2_O_2_ is schematically depicted. (**A)**. H_2_O_2_ emission was supported by pyruvate (10mM) and malate (2mM) to generate maximal rates and with ADP to assess H_2_O_2_ emission during OXPHOS. Maximal mitochondrial H_2_O_2_ along with the addition of ADP and an analysis of ADP’s attenuating effects as a percentage of maximal H_2_O_2_ was assessed in Early-Stage mice permeabilized muscle fiber bundles **(B-D)**. This was repeated in Late-Stage mice **(E-G).** ADP-stimulated respiration supported by pyruvate (5mM) and malate (2mM) generating NADH was also assessed in the absence (No Creatine) and presence (20mM Creatine) of creatine within permeabilized fiber bundles. Mitochondrial respiration was assessed at submaximal concentrations (25μM, 100μM and 500μM) (**H-K**). Protein contents of ETC subunits were also quantified within the TA **(L-N).** A two-way ANOVA was used for figures C, D, F, G, H-K. All data was analyzed using a one-way ANOVA and followed by a two-stage step-up method of Benjamini, Krieger and Yukutieli multiple comparisons test. Data that was not normally distributed was analyzed with a Kruskal-Wallis test followed by the same post-hoc analysis. Sham Control (Sham), Early-Stage Cancer Control (Early-EOC CON), Early-Stage Cancer SkQ1 (Early-EOC SkQ1), Late-Stage Cancer Control (Late-EOC CON), Late-Stage Cancer SkQ1 (Late-EOC SkQ1), electron transport chain (ETC), Arbitrary Units (AU). Results represent mean ± SD. n= 10-12. *β P*<0.05 Sham Vs Late-EOC CON; *γ P*<0.05 Sham Vs Late-EOC SkQ1; $ *P*<0.05 Late-EOC CON Vs Late-EOC SkQ1.

Mitochondrial respiration was also assessed by stimulating complex I with NADH generated by pyruvate and malate across a range of ADP concentrations to challenge mitochondria over a spectrum of metabolic demands. The ADP titrations were repeated without (No Creatine) and with creatine (20mM Creatine) in the assay to model the two main theoretical mechanisms of high energy phosphate shuttling from the mitochondria to the cytosol (16–18). See **SFigure7** for mechanistic modeling and theoretical framework partially adapted from our previous work (8). There were no differences in mitochondrial respiration in either creatine conditions within the Early-Stage time point **(Figure 4H and 4I).** This was consistent with pyruvate and malate in the absence of ADP (State II; reflective of proton leak through uncoupling pathways) or ADP-stimulated respiration further supported by glutamate (NADH) and succinate (Complex II stimulation via FADH_2_) **(SFigure 4A-4F)**. Within the Late-Stage timepoint, mitochondrial respiration was decreased in Late-EOC CON mice compared to Sham but unexpectedly further decreased in Late-EOC SkQ1 mice **(Figure 4J and 4K).** Interestingly, this effect of SkQ1 seemed to be specific to pyruvate and malate supported mitochondrial respiration as other substrates did not show a further decrease of mitochondrial respiration with SkQ1 treatment **(SFigure 4A-4F).** In addition, the ability of creatine to raise sub-maximal respiration was not different in all groups (analysis not shown). Last, none of the changes in mitochondrial respiration were explained by mitochondrial electron transport chain protein content markers given there were no changes at all timepoints and conditions **(Figure 4L-N).**

### SkQ1 administration in drinking water increases diaphragm mitochondrial complex I-stimulated H_2_O_2_ emission at Early- and Late-Stage cancer time points but attenuated cancer-induced reductions in Complex I protein content

Similar to the TA, we evaluated mitochondrial bioenergetics in the diaphragm to identify mechanisms of improved force production independent of atrophy. Complex I stimulated mH_2_O_2_ **(Figure 5A)** was completed in an identical paradigm as the TA. Within the Early-Stage timepoint, maximal mH_2_O_2_ and attenuations of mH_2_O_2_ with ADP were unchanged in Early-EOC CON and Early-EOC SkQ1 mice vs Sham **(Figure 5B and 5C)**. However, mH_2_O_2_ in the presence of ADP was increased in Early-EOC SkQ1 mice compared to Sham when expressed relative to maximal mH_2_O_2_ **(Figure 5D).** This unexpected finding suggests Complex I-supported mH_2_O_2_ increased with chronic SkQ1 treatment at Early-Stage cancer development due in part to less attenuation by ADP. Moreover, within the Late-Stage timepoint, maximal mH_2_O_2_ was unchanged but mH_2_O_2_ with ADP was increased in Late-EOC CON mice and further increased in Late-EOC SkQ1 mice **(Figure 5E and 5F)**. This trend remained when ADP’s attenuating effects were expressed as a percentage of maximal mH_2_O_2_ **(Figure 5G).** Normalizing maximal mH_2_O_2_ and mH_2_O_2_ in the presence of ADP to Complex I subunit protein content in **(Figure 5L & 5M)** does not change conclusions (analysis not shown) suggesting changes in mH_2_O_2_ were due to intrinsic properties independent of Complex I content. There were also no differences in the protein content of 4HNE adducts in Early- and Late-Stage mice vs Sham within the diaphragm **(SFigure 5B).** Collectively, this analysis suggests that late-stage ovarian cancer increases Complex I-supported mH_2_O_2_ and that diaphragm mitochondria appear to adapt to SkQ1 by increasing mH2O2 when assessed in vitro.

**Figure 5.**
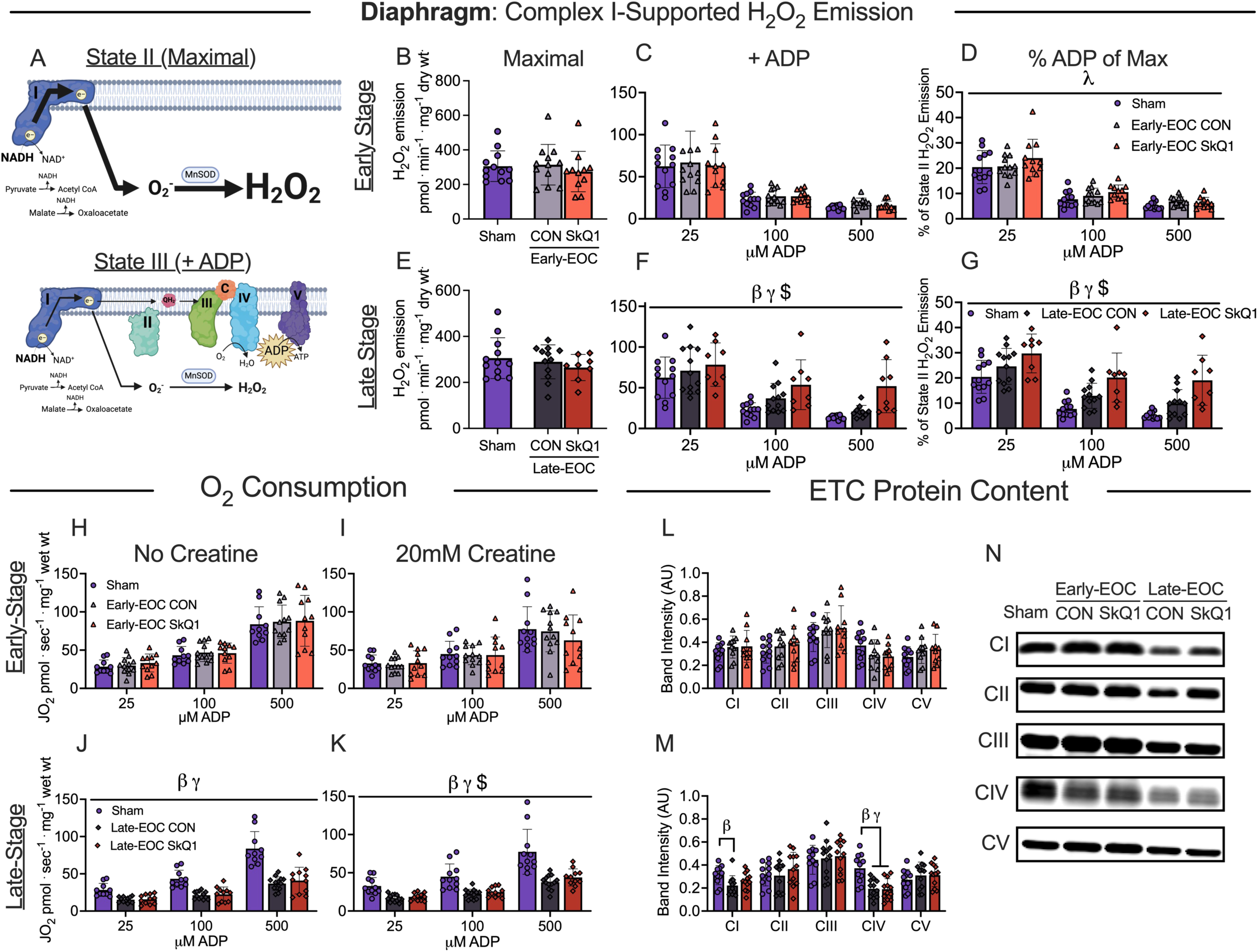
SkQ1 treatment in drinking water increases Late-Stage complex I stimulated H_2_O_2_ and, ADP-supported mitochondrial respiration in diaphragm muscle within a metastatic ovarian cancer cachexia mouse model. Complex I stimulated H_2_O_2_ is schematically depicted. (**A)**. H_2_O_2_ emission was supported by pyruvate (10mM) and malate (2mM) to generate maximal rates and with ADP to assess H_2_O_2_ emission during OXPHOS. Maximal mitochondrial H_2_O_2_ along with the addition of ADP and an analysis of ADP’s attenuating effects as a percentage of maximal H_2_O_2_ was assessed in Early-Stage mice permeabilized muscle fiber bundles **(B-D)**. This was repeated in Late-Stage mice **(E-G).** ADP-stimulated respiration supported by pyruvate (5mM) and malate (2mM) generating NADH was also assessed in the absence (No Creatine) and presence (20mM Creatine) of creatine within permeabilized fiber bundles. Mitochondrial respiration was assessed at submaximal concentrations (25μM, 100μM and 500μM) (**H-K**). Protein contents of ETC subunits were also quantified within the diaphragm **(L-N).** A two-way ANOVA was used for figures C, D, F, G, H-K. All data were analyzed using a one-way ANOVA and followed by a two-stage step-up method of Benjamini, Krieger and Yukutieli multiple comparisons test. Data that was not normally distributed was analyzed with a Kruskal-Wallis test followed by the same post-hoc analysis. Sham Control (Sham), Early-Stage Cancer Control (Early-EOC CON), Early-Stage Cancer SkQ1 (Early-EOC SkQ1), Late-Stage Cancer Control (Late-EOC CON), Late-Stage Cancer SkQ1 (Late-EOC SkQ1), electron transport chain (ETC), Arbitrary Units (AU). Results represent mean ± SD. n= 10-12. *β P*<0.05 Sham Vs Late-EOC CON; *γ P*<0.05 Sham Vs Late-EOC SkQ1; $ *P*<0.05 Late-EOC CON Vs Late-EOC SkQ1.

There were no differences in mitochondrial respiration in either creatine condition within the Early-Stage time point in the diaphragm **(Figure 5H and 5I).** This relationship was consistent across pyruvate & malate in the absence of ADP (State II), glutamate (State III) and succinate (State III) supported respiration **(SFigure 6A-F)**. ADP-stimulated pyruvate/malate supported respiration decreased in Late-EOC CON and Late-EOC SkQ1 mice compared to Sham and marginally increased in Late-EOC SkQ1 vs Late-EOC mice within the 20mM Creatine condition **(Figure 5J and 5K)**. This modest increase in mitochondrial respiration with SkQ1 was unique considering this pattern did not exist with other substrates in either creatine condition, albeit a slight improvement in respiration was also observed with succinate in the presence of creatine **(SFigure 6A-6F).** The ratio of respiration in the presence vs absence of creatine was greater than 1.0 in Sham, as expected, with no changes between groups (analysis not shown). The lower respiration in Late-EOC CON was associated with lower content of a Complex I subunit **(Figure 5L-5N)** whereas the increased respiration with SkQ1 was associated with possible preservation of Complex I subunit content as it was not different from Late-EOC CON although no differences from Sham were noted.

### SkQ1 administration in drinking water changes the phosphorylation status of pyruvate dehydrogenase (PDH) in the diaphragm

Phosphorylation of PDH by pyruvate dehydrogenase kinase (PDK) -4 inhibits PDH activity thereby lowering pyruvate oxidation (19) **(Figure 6A)**. Protein content of PDH, phosphorylation of PDH at the serine 293 site (P-PDH), ratio of P-PDH/PDH and PDK4 contents were unchanged within the TA in Early-Stage **(Figure 6B-6E)** and Late-Stage mice, with the exception of increased PDK-4 protein content in Late-EOC CON and Late-EOC SkQ1 mice **(Figure 6F-6I)**. Within the diaphragm in Early-Stage mice, PDH, P-PDH and P-PDH/PDH were unchanged compared to Sham **(Figure 6J-6L)** whereas PDK4 protein content increased in Early-EOC CON and Early-EOC SkQ1 mice compared to Sham **(Figure 6M).** In Late-Stage mice, diaphragm PDH protein content was lower in Late-EOC CON mice and Sham **(*P* = 0.06, Figure 6N),** while P-PDH was similar between all groups **(Figure 6O)**. The ratio of P-PDH/PDH increased in Late-EOC CON mice vs Sham which was prevented by SkQ1 **(*P* = 0.05, Figure 6P)** suggesting inhibitions in PDH activity during cancer may be prevented by SkQ1. This pattern matches the lower pyruvate-supported respiration in Late-EOC CON diaphragm with partial preservation by SkQ1 **(Figure 5K)**. There were no differences in PDK4 content within the diaphragm in Late-Stage mice **(Figure 6Q)** but this does not rule out changes in PDK activity given this enzyme is highly regulated by allosteric factors. See **Figure 6R and 6S** for representative blots.

**Figure 6.**
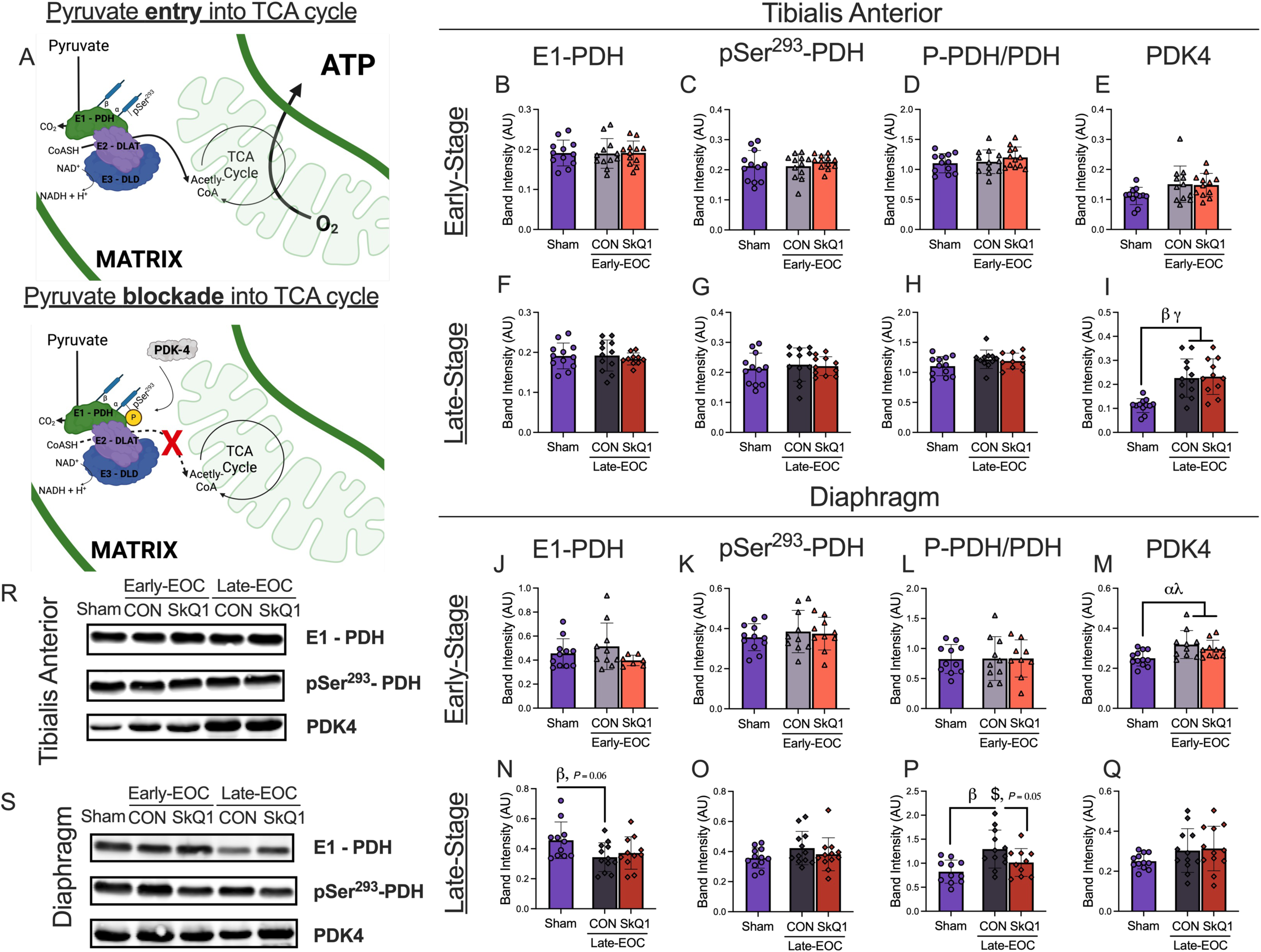
SkQ1 treatment in drinking water modulates phosphorylation status of pyruvate dehydrogenase in the diaphragm within a metastatic ovarian cancer cachexia mouse model. Theoretical schematic representation of pyruvate entry into mitochondria is depicted **(A)**. Protein contents of E1-PDH, P-PDH (Ser293), P-PDH/PDH and PDK4 were quantified in TA of Early- and Late-Stage mice (**B-I**, n=10-12). This was repeated in the diaphragm (**J-Q**, n=10-12). Representative blots (**R** and **S**). All data was analyzed using a one-way ANOVA and followed by a two-stage step-up method of Benjamini, Krieger and Yukutieli multiple comparisons test. Data that was not normally distributed was analyzed with a Kruskal-Wallis test followed by the same post-hoc analysis. Sham Control (Sham), Early-Stage Cancer Control (Early-EOC CON), Early-Stage Cancer SkQ1 (Early-EOC SkQ1), Late-Stage Cancer Control (Late-EOC CON), Late-Stage Cancer SkQ1 (Late-EOC SkQ1), Pyruvate dehydrogenase (PDH), Pyruvate dehydrogenase kinase (PDK), Arbitrary Units (AU); Dihydrolipoyl transacetylase (DLAT); Dihydrolipoamide dehydrogenase (DLD); carbon dioxide (CO_2_); Oxygen (O_2_); nicotinamide adenine dinucleotide (NADH); coenzyme A (CoASH). Results represent mean ± SD. *α P*<0.05 Sham Vs Early-EOC CON; *λ P*<0.05 Sham Vs Early-EOC SkQ1; *β P*<0.05 Sham Vs Late-EOC CON; *γ P*<0.05 Sham Vs Late-EOC SkQ1; $ *P*<0.05 Late-EOC CON Vs Late-EOC

### SkQ1 administration in the drinking water improves high-force cancer-induced muscle weakness and single fiber myoplasmic calcium release in the FDB

To further explore mechanisms of improved force production with SkQ1, we next evaluated *in-vitro* whole FDB muscle force production and single fiber intracellular free myoplasmic calcium concentrations ([Ca^2+^]i) in Early-Stage and Late-Stage mice. The FDB muscle was used as it is ideal for this experimental technique given the fibers are small/short (20). *In vitro* FDB force-frequency analyses was not different between all Early-Stage groups **(Figure 7A)**. However, single fiber [Ca^2+^]i were decreased in Early-EOC CON and Early-EOC SkQ1 mice compared to Sham **(Figure 7B)** suggesting disruptions to calcium handling occur early during ovarian cancer. While there was a greater tetanic [Ca^2+^]i that approached significance in Early-EOC SkQ1 mice vs Early-EOC CON across all frequencies **(*P* =0.08, Figure 7B)**, high-frequency stimulation analysis still demonstrates no muscle weakness occurred in Early-EOC CON despite decreases in tetanic [Ca^2+^]i within Early-EOC CON and also Early-EOC SkQ1 mice compared to Sham **(Figure 7C and 7D)**. At the late-stage timepoint, muscle force was unchanged in the FDB across all frequencies between each group **(Figure 7E)**. However, decreases in cancer-induced tetanic [Ca^2+^]i in Late-EOC CON vs Sham were partially prevented with SkQ1 treatment such that Late-EOC SkQ1 mice exhibited increases in tetanic [Ca^2+^]i compared to Late-EOC CON across all frequencies **(Figure 7F)**. At high-frequencies, FDB force production was lower in Late-EOC CON vs Sham mice but this was completely prevented by SkQ1 **(Figure 7G)**. Interestingly, this was related to a preservation of tetanic [Ca^2+^]i by SkQ1 compared to Late-EOC CON **(Figure 7H)** which suggests that SkQ1 counteracts cancer-induced decreases in single fiber sarcoplasmic reticulum (SR) calcium release which could be linked to improved force. Similarly, SkQ1 treatment partially preserved cancer-induced fatigue that occurred in early-stage cancer, but this was not related to increase single fiber SR calcium release throughout the fatigue protocol **(SFigure 8A and 8B).** SkQ1 treatment at Late-Stage also improved cancer-induced fatigue, however, this was coupled to improvements in single fiber SR calcium release throughout the fatigue protocol **(SFigure 8C and 8D).** Collectively, these results suggest improvements in muscle force production by SkQ1 are coupled to improvements in calcium handling.

**Figure 7.**
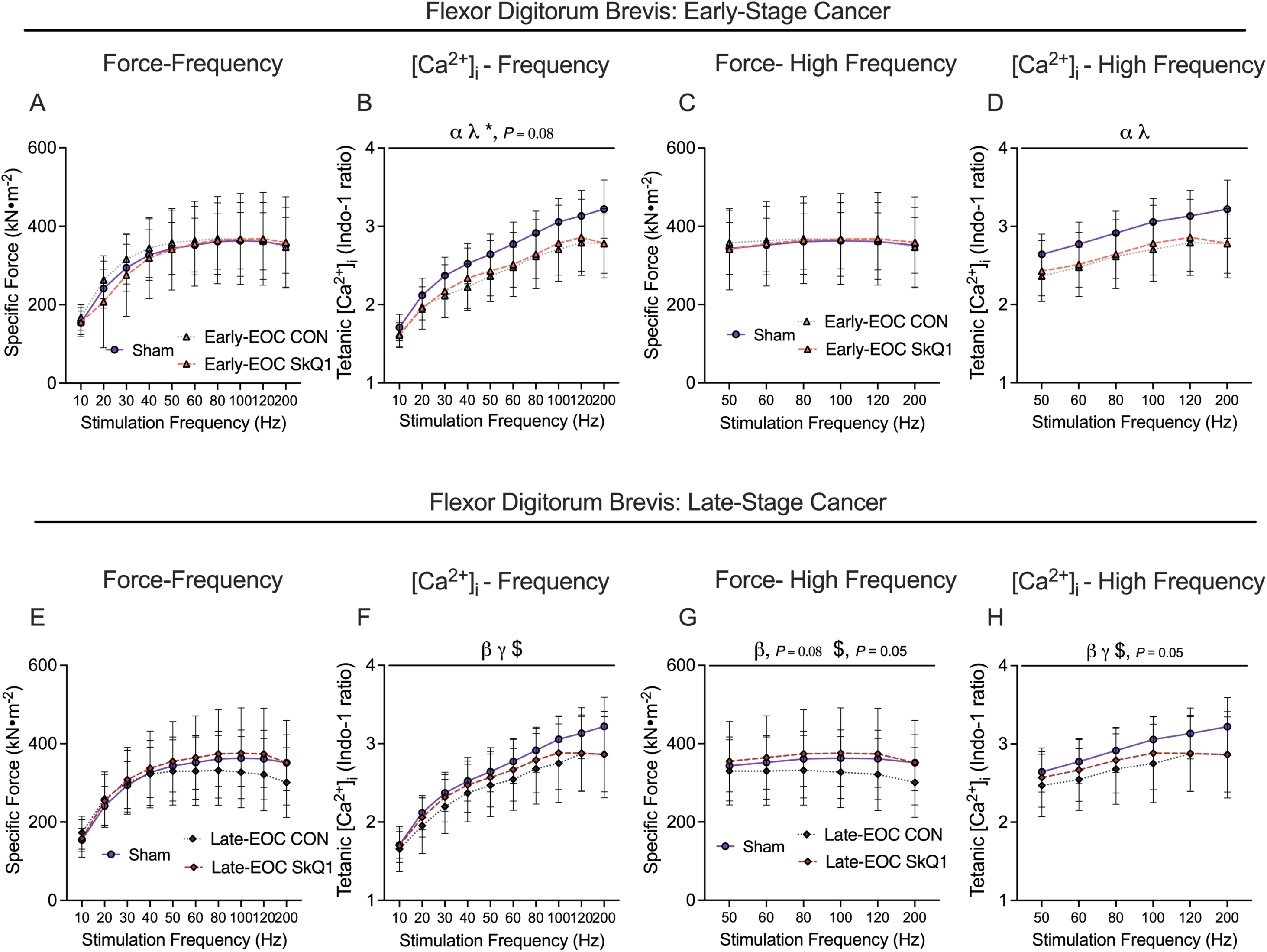
SkQ1 treatment in drinking water improves single fiber myoplasmic calcium release and force production in the flexor digitorum brevis within a metastatic ovarian cancer cachexia mouse model. In Early-Stage mice, in-vitro FDB force production was assessed using the force-frequency relationship (**A**, n =6-10). Myoplasmic calcium release as assessed using tetanic [Ca^2+^]_i_ across a range of frequencies (**B**, n=17-20), This analysis was repeated but only evaluating high-stimulation frequencies (**C, D**). All analysis was repeated in Late-Stage mice (**E-H**, n=8-17). A two-way ANOVA was used for all figures followed by a two-stage step-up method of Benjamini, Krieger and Yukutieli multiple comparisons test. [Ca^2+^]_i_ (intracellular calcium), Sham Control (Sham), Early-Stage Cancer Control (Early-EOC CON), Early-Stage Cancer SkQ1 (Early-EOC SkQ1), Late-Stage Cancer Control (Late-EOC CON), Late-Stage Cancer SkQ1 (Late-EOC SkQ1). Results represent mean ± SD. *α P*<0.05 Sham Vs Early-EOC CON; *λ P*<0.05 Sham Vs Early-EOC SkQ1; * *P*<0.05 Early-EOC CON Vs Early-EOC SkQ1; *β P*<0.05 Sham Vs Late-EOC CON; *γ P*<0.05 Sham Vs Late-EOC SkQ1; $ *P*<0.05 Late-EOC CON Vs Late-EOC SkQ1.

## Discussion

The recent discovery that reductions in muscle force can occur before the development of atrophy during cancer (7, 8) indicates that mechanisms beyond muscle size must contribute to weakness. Furthermore, the degree to which atrophy-independent causes of weakness occur in fibers following atrophy also remains unclear. Using a novel orthotopic, syngeneic, and immunocompetent mouse model of metastatic ovarian cancer recently reported by our group (8) this investigation demonstrates that the mitochondrial-targeted compound SkQ1 partially prevents weakness in the diaphragm before and after the development of atrophy without preventing atrophy itself. Likewise, SkQ1 partially prevented weakness in the TA without affecting atrophy which suggests that the contributions of mitochondrial stress to weakness during cancer is ubiquitous across muscle types. These discoveries indicate that mitochondria contribute to an atrophy-independent weakness during early and late stages of ovarian cancer progression and highlight the potential for mitochondrial-enhancing compounds to maintain muscle quality before and during cachexia.

The lack of effect of SkQ1 on survival in the present study does not detract from the importance of preserving muscle strength as an independent outcome during cancer given i) the findings serve as a basis for exploring the impact of preserving muscle strength with SkQ1 in cancer model combined with cancer therapy, ii) muscle quality is increasingly recognized as a predictor of survival in cancer patients undergoing cancer therapy (21), and iii) muscle strength predicts quality of life and reliance on health care systems.

Several notable relationships between mitochondrial metabolism, calcium handling and muscle quality were discovered as follows.

### SkQ1 administration exhibits muscle-specific effects on mitochondrial bioenergetics

Our results demonstrate that mitochondrial responses to SkQ1 are heterogenous between the TA and diaphragm in several ways. First, SkQ1 lowered Complex I-supported mH_2_O_2_ in the TA of Late-Stage mice specifically by enhancing the ability of ADP to suppress mH_2_O_2_. This discovery was made by designing *in vitro* assessments of mH_2_O_2_ in a way that recognizes the critical role of ADP as a regulator of reactive oxygen species generated by the electron transport chain. ADP stimulates proton re-entry through ATP synthase which lowers membrane potential and reduces the probability of premature electron slip onto oxygen and subsequent superoxide production prior to its dismutation to H_2_O_2_ (16). In this way, comparisons of mH_2_O_2_ in the absence vs presence of ADP *in vitro* revealed that mitochondria were less responsive to ADP during late-stage cancer and that SkQ1 partially prevented this apparent desensitization. While the precise mechanism of altered ADP-suppression of mH_2_O_2_ was not examined, the results direct new focus to the regulation of the phosphorylation system as possible targets of redox modifications, namely ADP/ATP cycling across the mitochondrial double membrane system (VDAC, ANT) or ATP synthase itself.

While this discovery during late-stage cancer links mH_2_O_2_ to muscle weakness but not atrophy in the TA, the prevention of weakness during early-stage cancer was not associated with altered mH_2_O_2_. This is difficult to explain for a number of reasons. First, it is not clear if SkQ1 was removed from mitochondria during sample preparation and whether SkQ1 must be present in the mitochondria to capture its effects during the *in vitro* assessments Likewise, it is possible that changes in response to SkQ1 treatment seen in Late-Stage TA may be due to adaptive reprogramming of the phosphorylation system that did not require the presence of SkQ1 during assessments. The latter consideration might also be relevant to why diaphragm mH_2_O_2_ was elevated in samples from SkQ1-treated mice. For example, it is not clear if mitochondria in the diaphragm relied on SkQ1 as a pseudo-‘crutch’ that led to an adaptive response of reduced intrinsic capacities to generate mH_2_O_2_. Furthermore, while we considered measuring SkQ1 in permeabilized fibers from drug-treated mice to determine if it was present during the assays, the tissue requirements far exceed the size of bundles, and we are not aware of an assay sensitive enough to measure its content in small samples. Nonetheless, the results clearly demonstrate that SkQ1 can partially prevent muscle weakness, but further work is required to perform comprehensive redox proteomics to determine if proteins regulating contractile function were modified by mitochondrial reactive oxygen species which is indeed warranted given SkQ1 is a well-established mitochondrial-targeted antioxidant and partially prevented weakness in this investigation.

Chronic SkQ1 treatment also decreased pyruvate oxidation in the TA despite modest increases in the diaphragm. These effects occurred despite no changes in the protein contents of subunits in the electron transport chain, which are stimulated during the *in vitro* assessments of respiration and therefore contribute to the measurement of respiration. Rather, the reduced diaphragm mitochondrial pyruvate oxidation in late-stage cancer, and its partial preservation by SkQ1, were linked to corresponding changes in PDH phosphorylation. However, while this was not observed in the diaphragm during early-stage cancer nor in the TA at either stage, allosteric control of PDH activity by acetyl CoA:CoA:, NADH:NAD, calcium and pyruvate itself (22) were not assessed which highlights the opportunity for further consideration of PDH activity as a potential site of altered metabolic regulation during cancer cachexia as proposed previously (23). This proposal, and the present data, offers possible considerations for future research to examine how reprogramming of carbohydrate oxidation may be linked to altered energy homeostasis or perhaps non-energy fates of glucose that have yet to be explored during cancer cachexia.

While the partial prevention of diaphragm weakness prior to atrophy is a unique finding of this investigation, the prevention of muscle weakness assessed by force frequency stimulation once atrophy occurs is consistent with a previous report with the mitochondrial cardiolipin-targeted peptide SS31 which also prevented diaphragm weakness in late-stage cancer in C26-ectopic tumour bearing mice (10). However, SS31 also prevented reductions in muscle CSA whereas this was not the case for SkQ1 in the present investigation. The reason for this discrepancy is unclear. While each drug has a different mechanism of action mediating reductions in mitochondrial reactive oxygen species and preserved oxidative phosphorylation, the different results may be more related to unique aspects of each cancer model due to cancer type, rate of cancer progression, presence of metastasis, or many other considerations. In this way, continued research into mitochondrial-enhancing compounds as potential agents to prevent atrophy vs atrophy-independent causes of weakness could be informed by considering how animal models capture the systemic stressors of cancer that occur in humans across time.

### SkQ1 administration improves force production potentially by improving calcium handling

We discovered ovarian cancer reduced tetanic [Ca^2+^]_i_ at the single fiber level and force at the whole fiber level within the FDB which were partially prevented by SkQ1 at high stimulation frequencies. This suggests that muscle weakness during ovarian cancer is caused - at least in part - by decreased calcium release during contraction, which is consistent with findings from other cancer models (24) but in a manner that is regulated to some effect by mitochondria. With this in mind, we attempted to immunoprecipitate ryanodine receptors (RyR1) for assessments of cysteine oxidation but were unsuccessful with the available tissue. Still, several lines of evidence warrant future consideration of calcium leak from RyR1 channels. For example, calcium leak through RyR1 is linked to muscle weakness during bone cancer (24) and ageing (25) due to RyR1 cysteine-nitrosylation and loss of callstabin1 – a channel stabilizing subunit (25). Likewise, aged MCAT mice (overexpression of the human catalase gene targeted to mitochondria) demonstrated less RyR1 oxidation, improved calcium release properties, and improved muscle force (26). These findings of RyR1 oxidation and muscle weakness may be unique to disease states given another study reported that *in vitro* administration of the mitochondrial-targeted antioxidant SS-31 to skeletal muscle from healthy mice restores fatigue-induced decreases in SR calcium release but does not improve force recovery (27). Collectively, future assessments of skeletal muscle RyR1 oxidation during cancer are warranted to determine whether a mitochondrial-SR redox signaling relay mediates muscle weakness hat can be modulated therapeutically with mitochondrial-targeted antioxidants.

### SkQ1 treatment attenuates mRNA expression of some atrogenes and cytokines

This investigation identified that SkQ1 lowers mRNA expression of *Atrogin* within the TA and *Il6* in addition to *Tnf* within the diaphragm. These findings are similar to the effects of SS-31 which attenuated C26 cancer-induced increases in mRNA expression of *Murf1*within the diaphragm and plasma levels of *Tnf* (10). Thus, mitochondrial-targeted antioxidants may be novel indirect ‘anti-inflammatories’ at least with regards to muscle-specific expression of gene programs regulating cytokines during cancer. However, the relationship between atrogenes and atrophy during cancer appears heterogeneous given we found SkQ1 prevented increases in mRNA of *Atrogin* in the TA during ovarian cancer with no changes in muscle fiber CSA. In a similar example, orthotopic breast cancer-induced reductions in soleus wet weight are not reversed in mice overexpressing the H_2_O_2_-specific antioxidant catalase in mitochondria. This demonstrates another model by which reducing mitochondrial ROS does not preserve muscle mass (28). However, SS-31 has been shown to attenuate mRNA of E3 ligases and preserve skeletal muscle CSA in the C26 model of cancer cachexia (10). While speculative, these differences could be due to the complexity of stressors in orthotopic and metastatic models of cancer cachexia (the present investigation) vs ectopic models. Furthermore, our findings might suggest other mechanisms govern skeletal muscle atrophy during ovarian cancer. For example, autophagy governed by Beclin1 and microtubule-associated protein 1 light chain 3B (LC3B) or apoptosis governed through caspase activation as been reported in other cancer models (29, 30) that were not measured in this study. However, such measures were not the focus of our questions regarding mitochondrial relationships to weakness. It is also possible that the introduction of ID8 cells at the selected ages (∼10 weeks for Late-Stage, ∼16 weeks for Early-Stage) disrupted growth processes to different degrees and distinct from atrophy processes. While growth processes were not assessed, the increased atrogenes in the Late-Stage group nonetheless point to a relationship with atrophy itself.

### Perspectives and Conclusions

These discoveries indicate that pharmacological targeting of mitochondria with plastoquinone-based compounds conjugated to lipophilic cations partially prevents muscle weakness without preventing atrophy in two distinct muscle types during ovarian cancer. While atrophy is a well-established cause of weakness, the findings provide a new perspective that skeletal muscle weakness can also be attributed to mitochondria in a manner that may be distinct from the molecular causes of atrophy itself **(Figure 8)**. In addition, the data indicates that mitochondrial-targeted therapy can treat the recently identified pre-cachexia weakness as well as weakness that occurs after the onset of atrophy. As such, the discoveries support the continued development of therapies to help preserve muscle function in addition to muscle mass during early and advanced stages of cancer. Furthermore, the preserved calcium release during contraction as seen with SkQ1 treatment highlights an exciting opportunity to investigate the mechanisms by which mitochondrial stress may regulate the sarcoplasmic reticulum in muscle during cancer. In pursuit of these and other possible mitochondrial-linked mechanisms, the potential for other types of mitochondrial-targeted compounds to treat both weakness and atrophy, as shown in previous work (10), underscores the importance of considering the specific mechanisms of actions of mitochondrial therapeutics as this growing paradigm of pharmacology continues to develop. Lastly, pre-cachexia weakness could represent a new biomarker of early-stage cancer that opens an opportunity for early therapeutic interventions for both cancer treatment and preservation of muscle function and mass.

**Figure 8.**
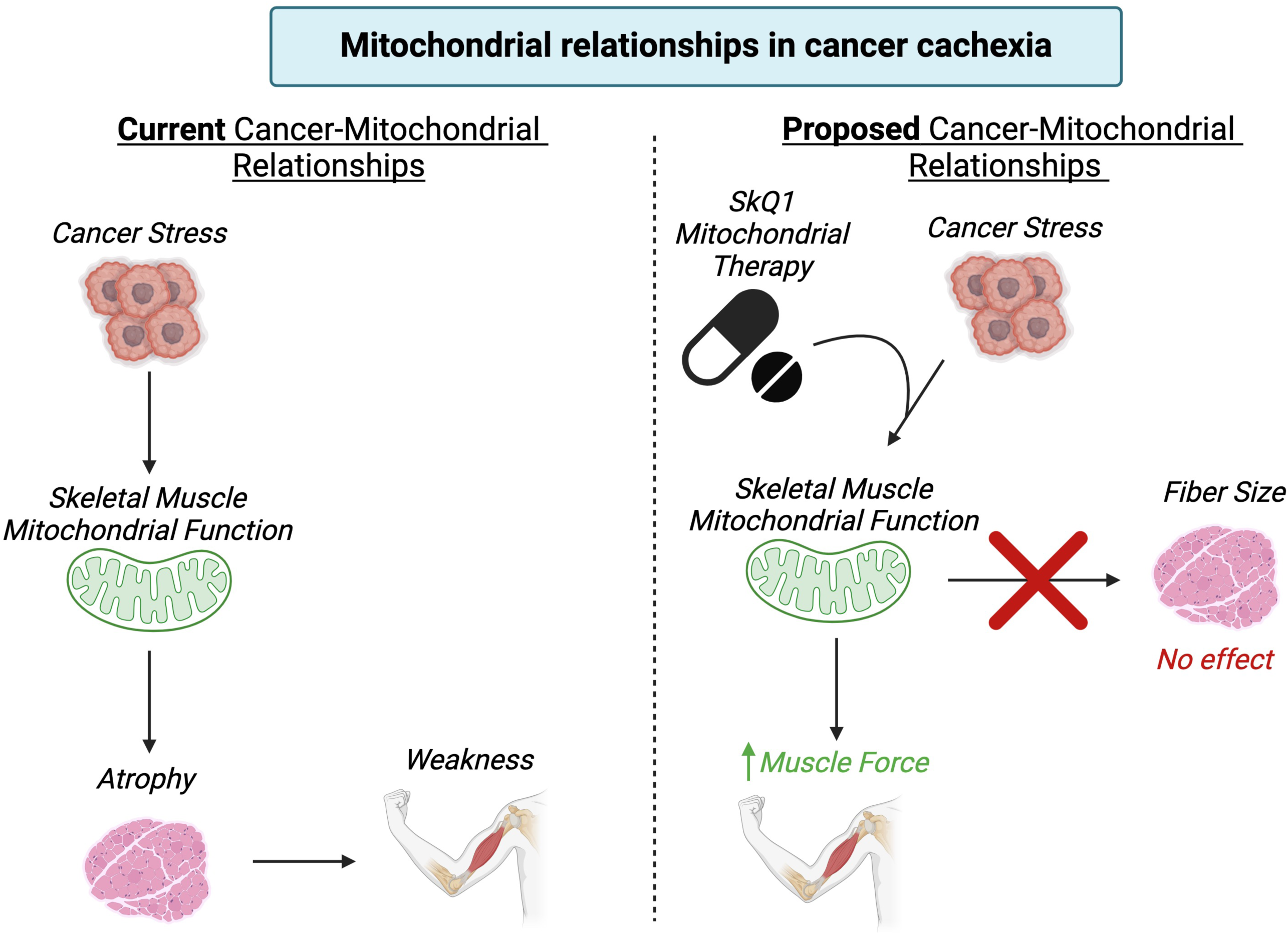
New proposed cancer-mitochondrial relationships after investigations with SkQ1 treatment in ovarian cancer cachexia. Current investigations in the relationships between mitochondrial function and cachexia suggest that cancer-induced alterations to mitochondrial function are linked to muscle atrophy which leads to muscle weakness (left). Using SkQ1 – a mitochondrial-targeted plastoquinone therapeutic – we identify a direct relationship between mitochondria and muscle force production/muscle weakness in cancer cachexia that are distinct from atrophy (right). Made with Biorender.

## Materials and Methods

### Animal Care and ID8 Inoculation

Female C57BL/6J mice (n=108) were ordered at 7-9 weeks of age from Charles River Laboratories. These mice were housed at the University of Guelph in accordance with the Canadian Council on Animal Care. Tumours were induced as described previously at the University of Guelph (8, 31–34). Briefly, ID8 cells (epithelial ovarian cancer cells; 1.0 x 10^6^ in 5μL) were injected under the left ovarian bursa of C57BL/6J mice generating an orthotopic model of ovarian cancer. Control mice were sham injected with equivalent volumes of sterile phosphate buffered saline (PBS). Two weeks after ID8 inoculation, mice were transported from University of Guelph to York University (AUP approval number 2021-04) where they were housed for the remainder of the study in accordance with the Canadian Council on Animal Care. All mice were provided access to irradiated Envigo 2918 chow and water *ad libitum*. Food weights were measured once a week from cages with group housed mice (4 mice per cage). Daily food intake was then estimated per cage by dividing the delta between initial food weight and final food weight by 7 (number of days) and 4 (number of mice). This same method was repeated for drinking water intake.

### Cancer Timepoints and Drug Incubations

Control mice (Sham; n =24) were sham injected at 10 weeks of age with PBS and aged to ∼23 weeks for end-point measures (13-15 weeks/91-105 days post-PBS injection). Four groups of mice received ID8 injections and divided into Early-Stage and Late-Stage subgroups. Each subgroup was further divided into those receiving SkQ1 or standard drinking eater (CON). Specifically, Early-Stage SkQ1 treated (Early-EOC SkQ1; n=12) and water treated (Early-EOC CON; n=12) mice were injected at 16-17 weeks and aged to ∼22-23 weeks for end-point measures (5-6 weeks/35 -46 days post ID8 injection). Late-Stage SkQ1 treated (Late-EOC SkQ1; n=30) and water treated (Late-EOC CON; n=30) mice were injected at 10 weeks and aged to ∼18-22 weeks for end-point measures (8-12 weeks/58-82 days post ID8 inoculation). A higher sample size was required for End-Stage mice given the unpredictability, heterogeneity and complications of cancer at this time. These time point ranges were partially informed by our previous work that reported similar ages for pre-atrophy weakness and atrophy in this mouse model (8). Mice in the Late-Stage group aged to the latest timepoint possible prior to presentation of combinations of the following criteria: 10% body weight loss, >20mL of ascitic volume collected during paracentesis, >5 ascites paracentesis taps completed, and/or subjective changes in behavioural patterns consistent with removal criteria as per animal care guidelines (self-isolation, ruffled fur/poor self-grooming and irregular gait) similar to our previous research(8). Mice in all groups received initial PBS or ID8 cells at slightly staggered age in order to ensure a sustainable rate of endpoint assessments given the requirement for fresh tissue for a number of experimental assays listed below.

SkQ1 (50 mg/ml) was purchased from Cayman Chemical (Item No 19891) in a solution of 1:1 ethanol:water. SkQ1 was administered in the drinking water such that all SkQ1-treated mice received 250nmol/kg body weight/day, as done previously (35–38). Mice were estimated to weigh 25g on average and drink 5 mL/day. Therefore, a 1.25nmol/mL concentration of SkQ1 was provided to all Early-SkQ1 and Late-SkQ1 mice from the day they received cancer injections. The estimated percentage of ethanol in the 1.25nmol/mL drinking water was 0.0008% as it was taken from the 50mg/mL 1:1 ethanol:water stock. Considering this was a very small percentage, Early-Veh and Late-Veh mice did not receive diluted ethanol in drinking water. SkQ1 contains a triphenylphosphonium (TPP) cation attached to the active quinone component that mediates the mitochondrial-targeted characteristic of the drug (13, 35). A TPP control was not included in this study given a previous investigation reported micromolar concentrations of TPP, which is higher than the drug dose used in the current study, did not interfere with cancer effects on atrophy nor explain the effects of MitoQ on grip strength, and a variety of biochemical and molecular measures in the C26 cancer model (11).

### Volitional Wheel Running

Forty-eight hours before euthanasia, a subset of mice were placed in individual cages with a 14 cm diameter running wheel and rotation counter (VDO m3 bike computer, Mountain Equipment Co-Op, Vancouver, Canada), as done previously (8, 39). Twenty-four hours later, running distance and time were recorded and mice were placed in separate caging with no running wheel. Muscle measurements were performed 24 hours thereafter. It was challenging to predict mouse survivability 48 hours in advance at the Late-Stage timepoint, therefore no volitional wheel running data was collected at this time. However, our previous work in this model has shown that mice do not typically run volitionally at late stages of ovarian cancer (8).

### Tissue Collection Procedure

Mice were anesthetized with isoflurane and euthanasia was performed by removal of the heart. All hindlimb muscles, inguinal subcutaneous fat and spleens were weighed and snap-frozen in liquid nitrogen and stored at -80^ο^C. Primary ovarian tumours were also collected by removing the ovary and tumour at the site of injection and carefully separating the tumour mass from the ovary mass and stored in liquid nitrogen. Hindlimb muscles were also embedded in optimal cutting temperature (OCT) medium and frozen (see section below). Tibialis anterior (TA) and diaphragm muscle were placed in BIOPS containing (in mM) 50 MES Hydrate, 7.23 K_2_EGTA, 2.77 CaK_2_EGTA, 20 imidazole, 0.5 dithiothreitol, 20 taurine, 5.77 ATP, 15 PCr, and 6.56 MgCl_2_·6 H_2_O (pH 7.1) to be prepared for mitochondrial bioenergetic assays. The flexor digitorum brevis (FBD) muscle was isolated and used for both *in-vitro* force production and enzymatic dissociation for myoplasmic calcium assessments (see sections below).

### In Situ Tibialis Anterior (TA) Force, In Vitro Diaphragm Force and In Vitro FDB Force

*In Situ* TA force production was partially adapted from previous literature (40, 41) and completed as done previously(8). Briefly, mice were anesthetized with isoflurane and the distal tendon of the TA was exposed by incision at the ankle. The distal tendon was tied with suture thread as close to the muscle attachment as possible. Once the knot was secured the distal tendon was severed. Small cuts were made up the lateral side of the TA to expose the muscle for needle electrode placement. The knot was tied to an Aurora Scientific 305C (Aurora Scientific, Aurora, ON, Canada) muscle lever arm with a hook. The foot of the mouse was secured with tape and the knee was immobilized with a needle and set screw with the length of the limb parallel to the direction of force. The two needle electrodes were placed in the gap of fascia between the TA and tibia to stimulate the common peroneal nerve (10-50 mA). The mouse was heated to maintain body temperature with a heating pad or heat lamp throughout force collection. Optimal resting length (L_o_) was determined using single twitches (pulse width=0.2ms) at 1 Hz stimulation frequency with 1 minute rest in between contractions to avoid fatigue. Once L_o_ was established, a ruler was used to determine TA length before the start of force-frequency collection (1, 10, 20, 30, 40, 50, 60, 80, 100, 120 and 200Hz with 1 minute rest in between contractions). Force production was normalized to the calculated CSA of the muscle strip (m/l*d) where m is the muscle mass, l is the length, and d is mammalian skeletal muscle density (1.06mg.mm^3^).

*In Vitro* diaphragm force production was completed as done previously(7). Briefly, the diaphragm strip was carefully sutured in Ringer’s solution (containing in mM: 121 NaCl, 5 KCl, 1.8 CaCl_2_, 0.5 MgCl_2_ 0.4 NaHPO_4_, 24 NaHCO_3_, 5.5 glucose and 0.1 EDTA; pH 7.3 oxygenated with 95% O_2_ and 5% CO_2_) such that the thread secured to the central tendon and ribs. The strip was then placed in an oxygenated bath filled with Ringer’s and maintained at 25^ο^C. The suture secured to the central tendon was then attached to the lever arm and the loop secured to the ribs was attached to the force transducer. The strip was situated between flanking platinum electrodes driven by biphasic stimulator (model 305C; Aurora Scientific Inc.). L_o_ determined using single twitches (pulse width=0.2ms) at 1 Hz stimulation frequency with 1 minute rest in between contractions to avoid fatigue. Once L_o_ was determined, the strip acclimatized for 30 minutes in the oxygenated bath. L_o_ was re-assessed and measured with a ruler and the start of the force-frequency protocol was initiated (1, 10, 20, 40, 60, 80, 100, 120, 140 and 200Hz with 1 minute rest in between contractions). Force production was normalized to cross-sectional area (CSA) of the muscle strip (m/(l*d)) where m is the muscle mass, l is the length, and d is mammalian skeletal muscle density (1.06mg.mm^3^).

*In Vitro* FDB force production was completed by attaching FDB tendons to 3D-printed clips and suture thread. The FDBs were then moved to a bath containing Tyrode solution oxygenated with 95% O_2_, 5% CO_2_ and pH ∼7.4 (model 6350*358, Aurora Scientific, Inc). One suture connected to the proximal end of FDB was attached to a lever arm, while the suture attached to the distal end of the FDB was attached to a force transducer. The FDB was placed between platinum electrodes connected to a stimulator (Aurora Scientific Integrated Muscle Test Controller). L_o_ was determined using twitches (pulse width = 0.2ms) at varying muscle lengths. Once L_o_ was determined, the strip acclimatized for 30 minutes in the oxygenated bath. Finally, force as a function of stimulation frequency was measured during 10 isometric contractions at varying stimulation frequencies (10, 20, 30, 40, 50, 60, 80, 100, 120, 200 Hz). Fatigue was tested during 120 repeated contractions every second (60Hz, 0.2ms pulse width, 0.3ms duration). Force production was normalized to the calculated CSA of the muscle strip (m/l*d) where m is the muscle mass, l is the length, and d is mammalian skeletal muscle density (1.06mg/mm^3^).

### Preparation of Permeabilized Muscle Fibers

The assessment of mitochondrial bioenergetics was performed as described previously in our publications(7, 8, 42–44). Briefly, the TA and diaphragm from the mouse was removed and placed in ice cold BIOPS. Muscle was separated gently along the longitudinal axis to form bundles that were treated with 40 μg/mL saponin in BIOPS on a rotor for 30 minutes at 4^ο^C. Following permeabilization the permeabilized muscle fiber bundles (PmFBs) for respiration were gently but quickly blotted and weighed in 1.5mL of tared prechilled BIOPS for normalization of respiratory assessments. The remaining PmFBs for mitochondrial H_2_O_2_ (mH_2_O_2_) were not weighed at this step as these data were normalized to fully recovered dry weights taken after the experiments. All PmFBs were then washed in Buffer Z on a rotator for 15 minutes at 4^ο^C to remove the cytoplasm. Buffer Z contained (in mM) 105 K-MES, 30 KCl, 10 KH_2_PO_4_, 5 MgCl_2_·6 H_2_O, 1 EGTA and 5mg/mL BSA (pH 7.1).

### Mitochondrial Respiration

High-resolution respirometry (O_2_ consumption) were conducted in 2 mL of respiration medium (Buffer Z) using the Oroboros Oxygraph-2k (Oroboros Instruments, Innsbruck, Austria) with stirring at 750 rpm at 37°C. Buffer Z contained 20 mM Cr to saturate mtCK and promote phosphate shuttling through mtCK or was kept void of Cr to prevent the activation of mtCK(45). All experiments were conducted in the presence of 5 μM blebbistatin (BLEB) in the assay media to prevent spontaneous contraction of PmFB, which has been shown to occur in response to ADP at 37°C that alters respiration rates(45). Complex I-supported respiration was stimulated using 5mM pyruvate and 2mM malate (NADH) followed by a titration of ADP concentrations from physiological ranges (25μM, 100μM(46)) to high submaximal (500μM) and saturating to stimulate maximal coupled respiration (5000μM in the presence of creatine and 7000μM in the absence of creatine). 10mM glutamate (further NADH generation) was added at the end of the ADP titration. Cytochrome *c* was then added to test mitochondrial outer membrane integrity. Experiments with low ADP-stimulated respiration (bundles that did not respond to ADP) with high cytochrome *c* responses (>15% increase in respiration) were removed from analysis. Lastly, 20mM Succinate (FADH_2_) was added to stimulate complex-II supported respiration. These protocols were designed to understand the regulation of respiration coupled to oxidative phosphorylation of ADP to ATP.

### Mitochondrial H_2_O_2_ emission (mH_2_O_2_)

mH_2_O_2_ was determined spectrofluorometrically (QuantaMaster 40, HORIBA Scientific) in PmfB placed in a quartz cuvette with continuous stirring at 37°C in 1 mL of Buffer Z supplemented with 10μM Amplex Ultra Red, 0.5 U/ml horseradish peroxidase, 1mM EGTA, 40 U/ml Cu/Zn-SOD1, 5μM BLEB and 20mM Cr to saturate mtCK. No comparisons were made to PmFB in the absence of creatine due to tissue limitations. State II mH_2_O_2_ (maximal emission in the absence of ADP) was induced using the Complex I-supporting substrates (NADH) pyruvate (10mM) and malate (2mM) to generate forward electron transfer (FET)-supported electron slip at Complex I (47) as described previously(39). These PmFBs were incubated with 35 μM CDNB during the 30-minute permeabilization to deplete glutathione and allow for detectable rates of mH_2_O_2_. Following the induction of State II mH_2_O_2_, a titration of ADP was employed to progressively attenuate mH_2_O_2_ as it occurs when membrane potential declines during oxidative phosphorylation(16). After the experiments, the fibers were lyophilized in a freeze-dryer (Labconco, Kansas City, MO, USA) for >4 hours and weighed on a microbalance (Sartorius Cubis Microbalance, Gottingen Germany). The rate of mH_2_O_2_ was calculated from the slope (F/min) using a standard curve established with the same reaction conditions and normalized to fiber bundle dry weight.

### [Ca^2+^]_i_: Enzymatic Dissociation of Intact Muscle Fibers

FDB muscles were dissected at room temperature in experimental Tyrode solution consisting of 121mM NaCl, 5.0mM KCl, 1.8mM CaCl_2_, 0.5mM MgCl_2_, 0.4mM NaH_2_PO_4_, 24mM NaHCO_3_, 0.1mM EDTA, and 5.5mM glucose. Glass-bottomed Petri dishes with a diameter of 35 mm were placed on a sterilized incubator tray, coated with laminin (Cat. CB-40232, Thermo Fisher Scientific, Waltham, MA, USA) and left at room temperature for ∼2 hours.

Isolated FDBs were transferred to a multiwell plate containing 3 mL of: Dulbecco’s modified Eagle’s medium (DMEM)/nutrient mixture F-12, pH 7.4 (Cat. 12800017, Thermo Fisher Scientific, Waltham, MA, USA); 20% heat-inactivated newborn calf serum (Cat. 16010159, Gibco, New Zealand); antibiotic antimycotic solution (6µL/mL; Cat. no. P4333 Sigma-Aldrich, St Louis, MO, USA); 0.6% collagenase Type I (Cat. SCR103, EMD Millipore Corp., Burlington, MA, USA), and 0.01% sodium bicarbonate. The multiwell plate was then incubated at 37°C in a water-saturated incubator for 2-3 hours. Meanwhile, glass bottom petri dishes (35 mm) were placed on a sterlized incubator tray, coated with laminin (Cat. CB-4032, Thermo Fisher Scientific, Waltham, MA, USA) and left at room temperature for ∼2 hours. Afterwards, FDB muscles were transferred into 1 mL of fresh DMEM/F-12 with 20% newborn calf serum and antibiotic antimycotic solution and carefully mechanically dissociated via pipetting the solution ∼30 times. The glass-bottomed Petri dishes coated with laminin for 2 hours were then washed with PBS and loaded with 165 µL of DMEM-F-12 solution containing the muscle fibers. The dishes were left for ∼45 minutes at room temperature to allow muscle fibers to adhere to the laminin coated glass. Afterwards, 3 mL of DMEM/F-12 with 20% newborn calf serum and antibiotic antimycotic solution were added. The petri dishes were then placed into the incubator and stored at 37 °C overnight and used the following day.

### [Ca^2+^]_i_: Dye Loading

The petri dishes containing the dissociated FDB muscle fibers were then loaded with [3.5μM] of indo-1 AM (Cat. I1223, Invitrogen) and placed in a water saturated incubator (37 °C) for 30 minutes. Dishes were then moved to a dark room and placed into a custom-made 3D printed chamber and mounted on an inverted microscope (Nikon Eclipse Ts2R-FL, Tokyo, Japan) where the remainder of the experiment was conducted at room temperature. The dishes were then washed for 30 minutes with a combination of experimental Tyrode solution, 0.5% newborn calf serum and bubbled with a mixture of 95% O_2_–5% CO_2_ (∼7.4 pH). Electrical stimulation was performed using the custom 3D printed chamber with two platinum electrodes (1cm apart) and controlled by an electrical stimulator (Model 701C, Aurora Scientific, Aurora, ON, Canada) via Aurora Scientific 600A software (Aurora Scientific, Aurora, ON, Canada). To assess [Ca^2+^]_i_, Indo-1AM (Cat. I1223, Invitrogen, Eugene, OR, USA) dye was excited at 346 ± 5 nm and the two emission wavelengths were recorded by two photomultipliers at 405±5 nm and 495±5 nm (Horiba Ratiomaster, London, ON, Canada) as done previously (20, 48). Felix32 software (Horiba, London, ON, Canada) was used to acquire [Ca^2+^]_i_ readings (resting and tetanic values).

### [Ca^2+^]_i_-frequency assessment and fatigue protocol

[Ca^2+^]_i_-frequency assessment was partially adapted from previous literature (49). The [Ca^2+^]_i_-frequency relationship was assessed by stimulating the fibers with 300ms tetani (0.2ms pulse duration) at progressively increasing stimulation frequencies of 1-200Hz (1, 10, 20, 30, 40, 50, 60, 80, 100, 120, 200Hz) with 1-minute intervals of rest between each tetanus. Fatigue was tested during 120 repeated contractions every second (60Hz, 0.2ms pulse width, 300ms duration).

### Sectioning & Immunofluorescence

TA and diaphragm muscle samples were first dried on a Kim wipe for ∼30 seconds to avoid freeze fracture. Muscles were then put on a longitudinally cut pipette tip and generously embedded in OCT medium (Thermo Fisher Scientific). Samples were then put in liquid nitrogen-chilled 2-methylbutane (ice visible on bottom of steel bucket, ∼3 minutes of chilling) for >60 seconds and transferred to Eppendorf tubes where they were kept at -80^ο^C for long term storage. These muscles were then sectioned into 10μm sections with a cryostat (HM525 NX, Thermo Fisher Scientific) maintained at -20^ο^C on Fisherbrand Superfrost Plus slides (Thermo Fisher Scientific). Immunofluorescent analysis of myosin heavy chain (MHC) expression was completed as previously described(50). Images were taken using a fluorescence slide scanner at wavelengths 350nm (blue), 488nm (green) and 555nm (red) (Zeiss AxioScan) at 20x magnification. Sections were analyzed using the ImageJ extension on QuPath-0.4.0 software while blinded to the groups. Fibers that did not fluoresce were considered IIx fibers. A total of 25-50 fibers were then randomly selected and measured. Type I fibers were extremely low in abundance in the TA and thus were not analyzed(50).

### RNA Isolation and Rt-PCR

RNA isolation was performed as previously described(51). TA and diaphragm samples were lysed using TRIzol reagent (Invitrogen) and RNA was separated to an aqueous phase using chloroform. The aqueous layer containing RNA was then mixed with isopropanol and loaded to Aurum Total RNA Mini Kit columns (Bio-Rad, Mississauga, ON, Canada). Total RNA was then extracted according to the manufacturer’s instructions. RNA was quantified spectrophotometrically using the NanoDrop attachment for the Varioskan LUX Multimode Microplate reader (Thermo Scientific). Reverse transcription of RNA into cDNA was performed by M-MLV reverse transcriptase and oligo(dT) primers (Qiagen, Toronto, ON, Canada). cDNA was then amplified using a CFX384 Touch Real-Time PCR Detection Systems (Bio-Rad) with a SYBR Green master mix and specific primers **(STable 1)**. Gene expression was normalized to β-actin (Actb) and relative differences were determined using the ΔΔCt method. Values are presented as fold changes relative to the control group.

### Western Blotting

A frozen piece of TA and diaphragm from each animal was homogenized in a plastic microcentrifuge tube with a polytron homogenizer in ice-cold buffer containing (in mM) 20 Tris/HCl, 150 NaCl, 1 EDTA, 1 EGTA, 2.5 Na_4_O_7_P_2_, and 1 Na_3_VO_4_ and 1% Triton X-100 with PhosSTOP inhibitor tablet (Roche; 4906845001) and protease inhibitor cocktail (Sigma Aldrich; P8340) (pH7.0) as published previously(7, 52). Protein concentrations were determined using a bicinchoninic acid assay (life Technologies, Thermo Fisher Scientific). Denatured and reduced protein (15-40 μg) was subjected to 10%-12% gradient SDS-PAGE followed by transfer to low-fluorescence polyvinylidene difluoride membrane. Membranes were blocked with Odyssey Blocking Buffer (Li-COR) and immunoblotted overnight (4^ο^C) with antibodies specific to each protein. A commercially available monoclonal antibody was used to detect electron transport chain proteins (rodent OXPHOS Cocktail, ab110413; Abcam, Cambridge, UK, 1:500 dilution), including V-ATP5A (55kDa), III-UQCRC2 (48kDa), IV-MTCO1 (40kDa), II-SDHB (30 kDa), and I-NDUFB8 (20 kDa). A commercially available monoclonal antibody was used to detect PDH (Abcam Anti-PDHA1 ab110330, 1:500 dilution), polyclonal antibody for P-PDH (P-PDHα1 (Ser293) CST #31866, 1:250 dilution), monoclonal antibody for PDK4 (Abcam Anti-PDK4 ab214938, 1:500 dilution) and monoclonal antibody for 4-HNE (Abcam HNEJ-2 ab48506, 1:500 dilution).

After overnight incubation in primary antibodies, membranes were washed 3 times for 5 minutes in TBS-Tween and incubated for 1 hour at room temperature with the corresponding infrared fluorescent secondary antibody (LI-COR IRDye 680LT 925-68021 Goat anti-Rabbit, 1:20,000; 680RD 925-68070 Goat anti-Mouse, 1:20,000). Images were then normalized to a whole membrane Amido Black total protein stain (A8181, Sigma).

### Statistics

Results are expressed as mean ± SD. The level of significance was established at *p* <0.05 for all statistics. The D’Agostino-Pearson omnibus normality test was first performed to determine whether data resembled a Gaussian distribution, and all data were subject to the ROUT test (Q=0.5%) to identify and exclude outliers which was a rare occurrence. When data fit normal distributions, one way and two-way ANOVAs were performed. When data did not fit a Gaussian distribution for analysis with one independent variable, the Kruskal-Wallis test was used. Moreover, when data did not fit a Gaussian distribution for analysis with two independent variables, data was first log transformed then analyzed using a two-way ANOVA (See **SFigure 9** for log transformed analysis of **Figure 4F**, **4J**, **4K and 5F**) but data was still presented in the Results as non-transformed data. Respective statistical tests are provided in figure legends. When significance was observed with an ANOVA, post-hoc analyses were performed with a two-stage set-up method of Benjamini, Krieger and Yekutieli for controlling false discovery rate (FDR) for multiple-group comparisons. With this method, all reported *p* values are FDR adjusted (traditionally termed “q”). All statistical analyses were performed in GraphPad Prism 10 (La Jolla, CA, USA).

The purpose of this study was to evaluate the effect of SkQ1 at both the Early-Stage and Late-Stage time points. Consequently, we have chosen to separate our statistical analysis by completing separate tests on Early- and Late-Stage timepoints but compared to a shared control group to reduce the number of mice used in this study in accordance with the Animal Care Committee. We did not analyze the impact of cancer over time (Early-Veh vs Late-Veh comparisons) as this analysis has already been completed by our group in the past (8).

## Study Approval

All experiments and procedures were approved by the Animal Care Committee at York University and the University of Guelph in accordance with the Canadian Council on Animal Care.

## Data availability

data are available upon reasonable request submitted to the corresponding author.

## Author contributions

L.J.D., S.K., S.T., J.A.S., N.P.G., A.J.C., J.P., and C.G.R.P. contributed to the rationale and study design. L.J.D., S.K., L.D.F., N.J.A., M.C.G., S.G., A.N.B., B.A.M., B.G., S.L., and C.A. conducted all experiments and/or analyzed all data. L.JD., and C.G.R.P performed project administration and wrote the manuscript. Funding was provided by C.G.R.P and J.A.S. All authors have approved the final version of the manuscript and agree to be accountable for all aspects of the work. All persons designated as authors qualify for authorship, and all those who qualify for authorship are listed.

## Acknowledgements

Funding was provided to CGRP by the National Science and Engineering Research Council (NSERC, 436138-2013 and 2019-06687) and an Ontario Early Research Award (2017-0351) with infrastructure supported by the Canada Foundation for Innovation, the Ontario Research Fund, and the James H. Cummings Foundation. LJD was supported by NSERC CGS-D scholarship. SK was supported by NSERC CGS-M scholarship. SG and AKT were supported by Ontario Graduate Scholarship (OGS). KAM was supported by National Institute of Health (NIH) R01 AG080047. JS and JP were supported by NSERC and Canadian Institute of Health Research (CIHR).

## Conflict of Interest

The authors have declared that no conflict of interest exists.

## Notes

### Competing Interest Statement

The authors have declared no competing interest.

