## Supplemental Figure 1 for "Mitochondrial-targeted plastoquinone therapy ameliorates early onset muscle weakness that precedes ovarian cancer cachexia in mice"

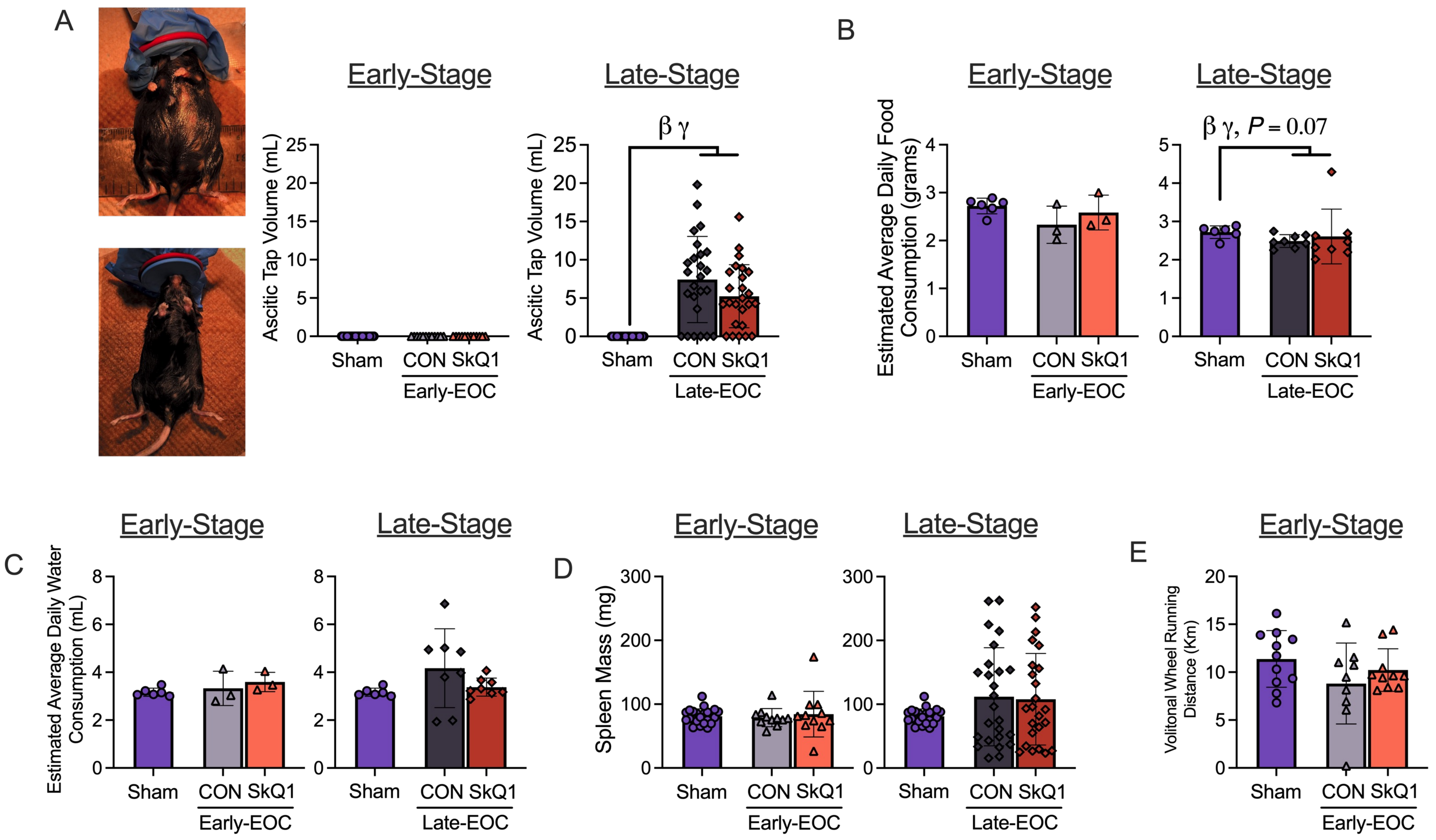


**SFigure 1. Effects of SkQ1 on ascitic volume, estimated daily food intake, estimated daily water intake, spleen mass and volitional wheel running.** Abdominal ascites was drained as required (**A**, n=24-25). Daily food intake was estimated as food weights were taken once a week and then divided by the number of mice in the cage and days since last food weight (**B**, n=3-8). This was repeated for estimated average daily water intake (**C**, n=3-8). Spleen mass (**D**, n=12-25) and volitional wheel running after 24 hours of exposure to a running wheel was also analyzed (**E**, n=9-12). All data was analyzed using a one-way ANOVA and followed by a two-stage step-up method of Benjamini, Krieger and Yukutieli multiple comparisons test. Data that was not normally distributed was analyzed with a Kruskal-Wallis test followed by the same post-hoc analysis. Sham Control (Sham), Early-Stage Cancer Control (Early-EOC CON), Early-Stage Cancer SkQ1 (Early-EOC SkQ1), Late-Stage Cancer Control (Late-EOC CON), Late-Stage Cancer SkQ1 (Late-EOC SkQ1). Results represent mean ± SD. $\beta$ $P<0.05$ Sham Vs Late-EOC CON; $\gamma$ $P<0.05$ Sham Vs Late-EOC SkQ1.
