## Supplemental Figure 2 for "Mitochondrial-targeted plastoquinone therapy ameliorates early onset muscle weakness that precedes ovarian cancer cachexia in mice"

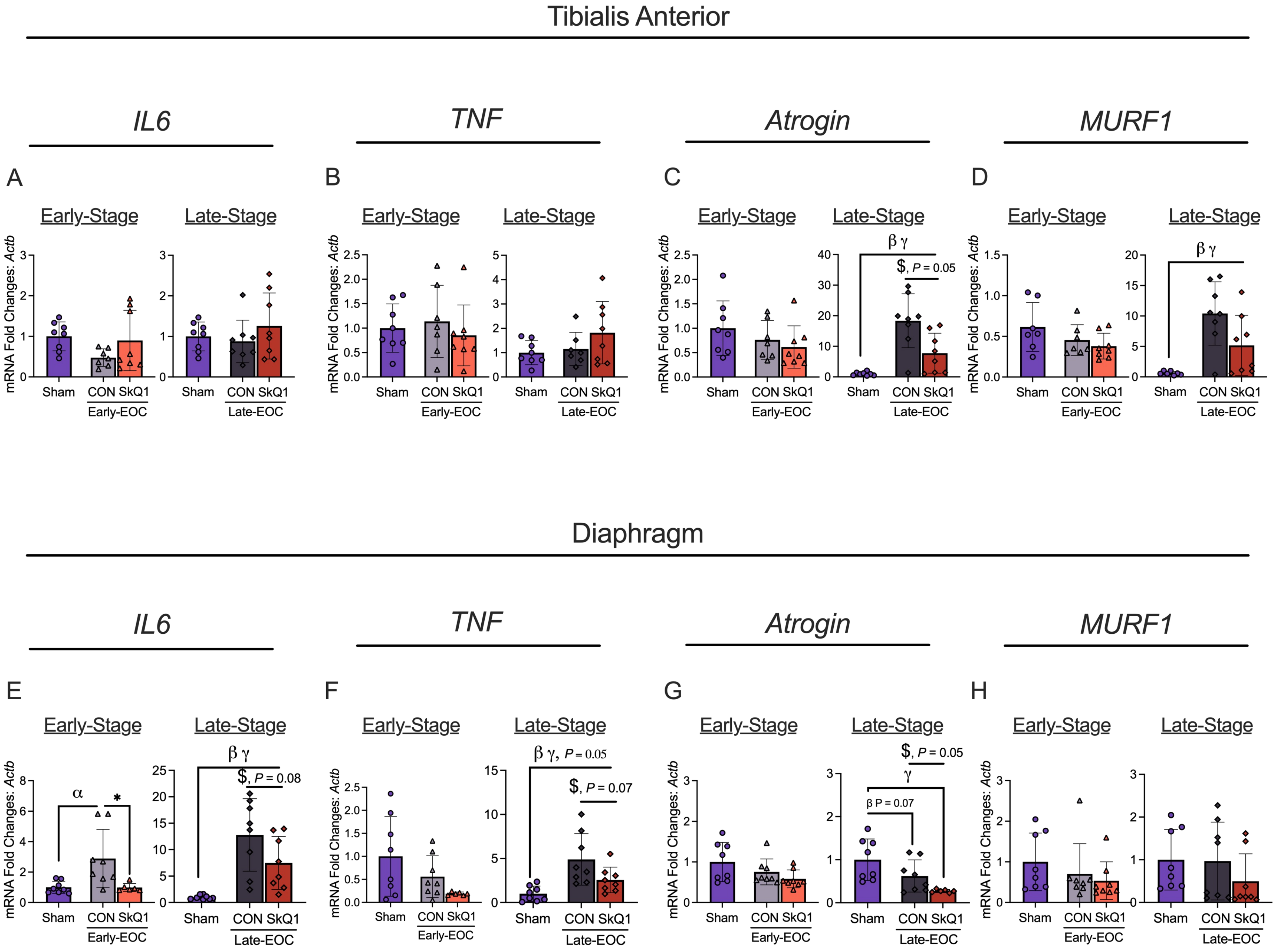


**SFigure 2. SkQ1 treatment in drinking water reduces Atrogin mRNA expression in tibialis anterior at Late-Stage tumour development and and IL-6 /TNF mRNA expression in the diaphragm within a metastatic ovarian cancer cachexia mouse model.** Analysis of mRNA expression fold changes in IL6, TNF, Atrogin and MURF1 was completed in the TA in Early-Stage and Late-Stage mice (**A-D**, n=8). This was repeated in the diaphragm (**E-H**, n=8). All data was analyzed using a one-way ANOVA and followed by a two-stage step-up method of Benjamini, Krieger and Yukutieli multiple comparisons test. Data that was not normally distributed was analyzed with a Kruskal-Wallis test followed by the same post-hoc analysis. Sham Control (Sham), Early-Stage Cancer Control (Early-EOC CON), Early-Stage Cancer SkQ1 (Early-EOC SkQ1), Late-Stage Cancer Control (Late-EOC CON), Late-Stage Cancer SkQ1 (Late-EOC SkQ1), *IL6* (Interleukin-6), *TNF* (Tumour necrosis factor), *MURF1* (RING-finger protein-1), TA (tibialis anterior), *actb* (Beta actin). Results represent mean ± SD. $\alpha$ $P$<0.05 Sham Vs Early-EOC CON; $\beta$ $P$<0.05 Sham Vs Late-EOC CON; $\gamma$ $P$<0.05 Sham Vs Late-EOC SkQ1, * $P$<0.05 Early-EOC CON Vs Early-EOC SkQ1, $ $P$<0.05 Late-EOC CON Vs Late-EOC SkQ1.
