## Supplemental Figure 3 for "Mitochondrial-targeted plastoquinone therapy ameliorates early onset muscle weakness that precedes ovarian cancer cachexia in mice"

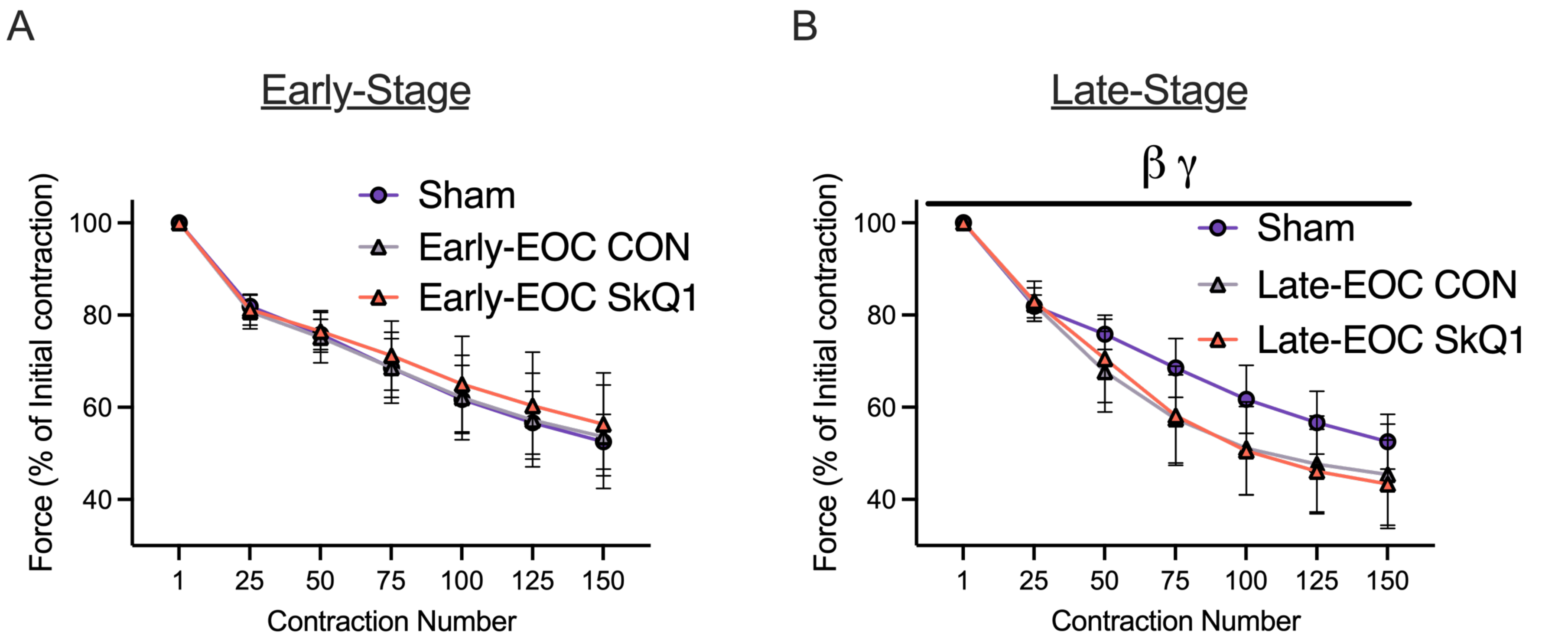


**SFigure 3. Analysis of force production within the diaphragm during a fatigue protocol.** In-Vitro diaphragm fatigue was completed (70Hz for 350ms every 2s for 150 contractions) to assess the effects of SkQ1 on fatiguability in Early-Stage and Late-Stage mice **(A, B)**. All data was analyzed using a two-way ANOVA and followed by a two-stage step-up method of Benjamini, Krieger and Yukutieli multiple comparisons test. Sham Control (Sham), Early-Stage Cancer Control (Early-EOC CON), Early-Stage Cancer SkQ1 (Early-EOC SkQ1), Late-Stage Cancer Control (Late-EOC CON), Late-Stage Cancer SkQ1 (Late-EOC SkQ1). Results represent mean ± SD. n=8-11. ; $\beta$ $P<0.05$ Sham Vs Late-EOC CON; $\gamma$ $P<0.05$ Sham Vs Late-EOC SkQ1.
