## Supplemental Figure 5 for "Mitochondrial-targeted plastoquinone therapy ameliorates early onset muscle weakness that precedes ovarian cancer cachexia in mice"

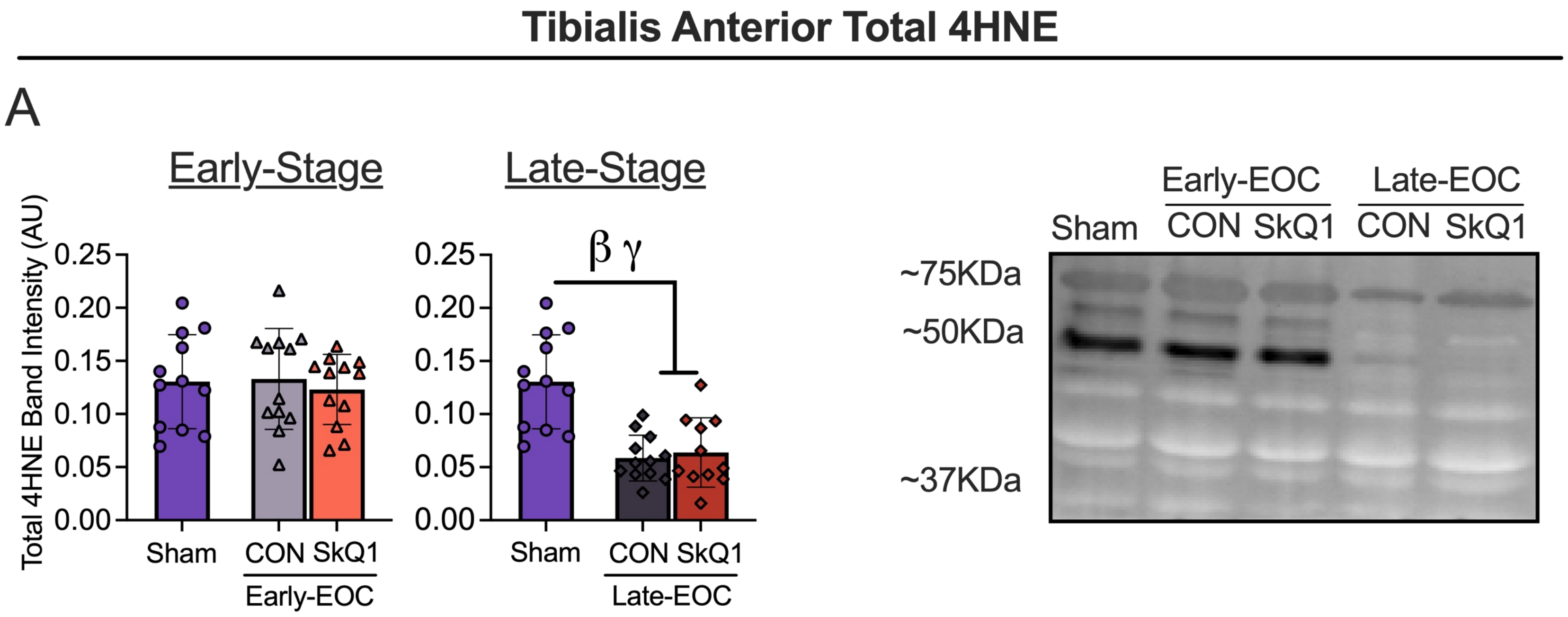


**SFigure 5. Quantification of 4HNE Western Blot Analysis.** Western blot assessment of 4HNE protein adduct formation were analyzed in the TA (**A;** n=11-12) and diaphragm (**B;** n=10-12). All data was analyzed using a one-way ANOVA and followed by a two-stage step-up method of Benjamini, Krieger and Yukutieli multiple comparisons test. Data that was not normally distributed was analyzed with a Kruskal-Wallis test followed by the same post-hoc analysis Sham Control (Sham), Early-Stage Cancer Control (Early-EOC CON), Early-Stage Cancer SkQ1 (Early-EOC SkQ1), Late-Stage Cancer Control (Late-EOC CON), Late-Stage Cancer SkQ1 (Late-EOC SkQ1), Arbitrary Units (AU). Results represent mean ± SD. n=10-12. $\beta$ $P<0.05$ ; $\beta$ $P<0.05$ Sham Vs Late-EOC CON; $\gamma$ $P<0.05$ Sham Vs Late-EOC SkQ1.

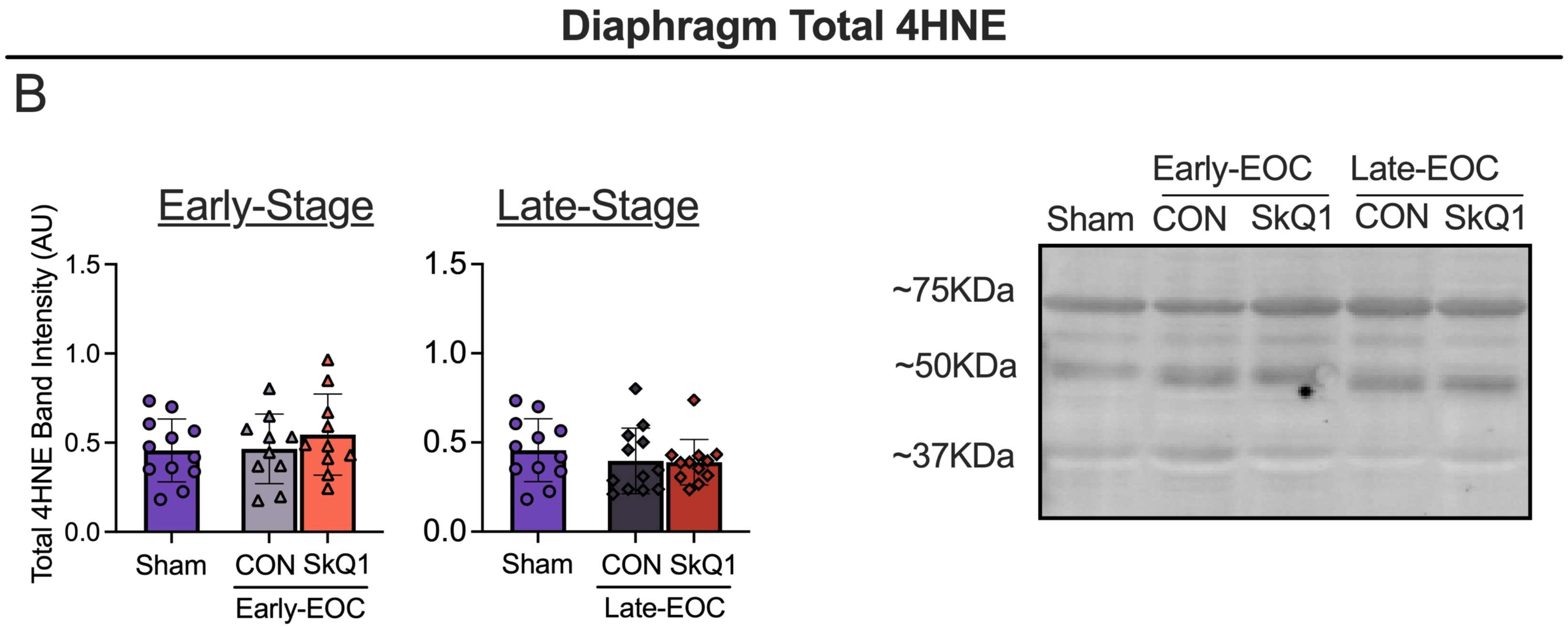
