## Supplemental Figure 6 for "Mitochondrial-targeted plastoquinone therapy ameliorates early onset muscle weakness that precedes ovarian cancer cachexia in mice"

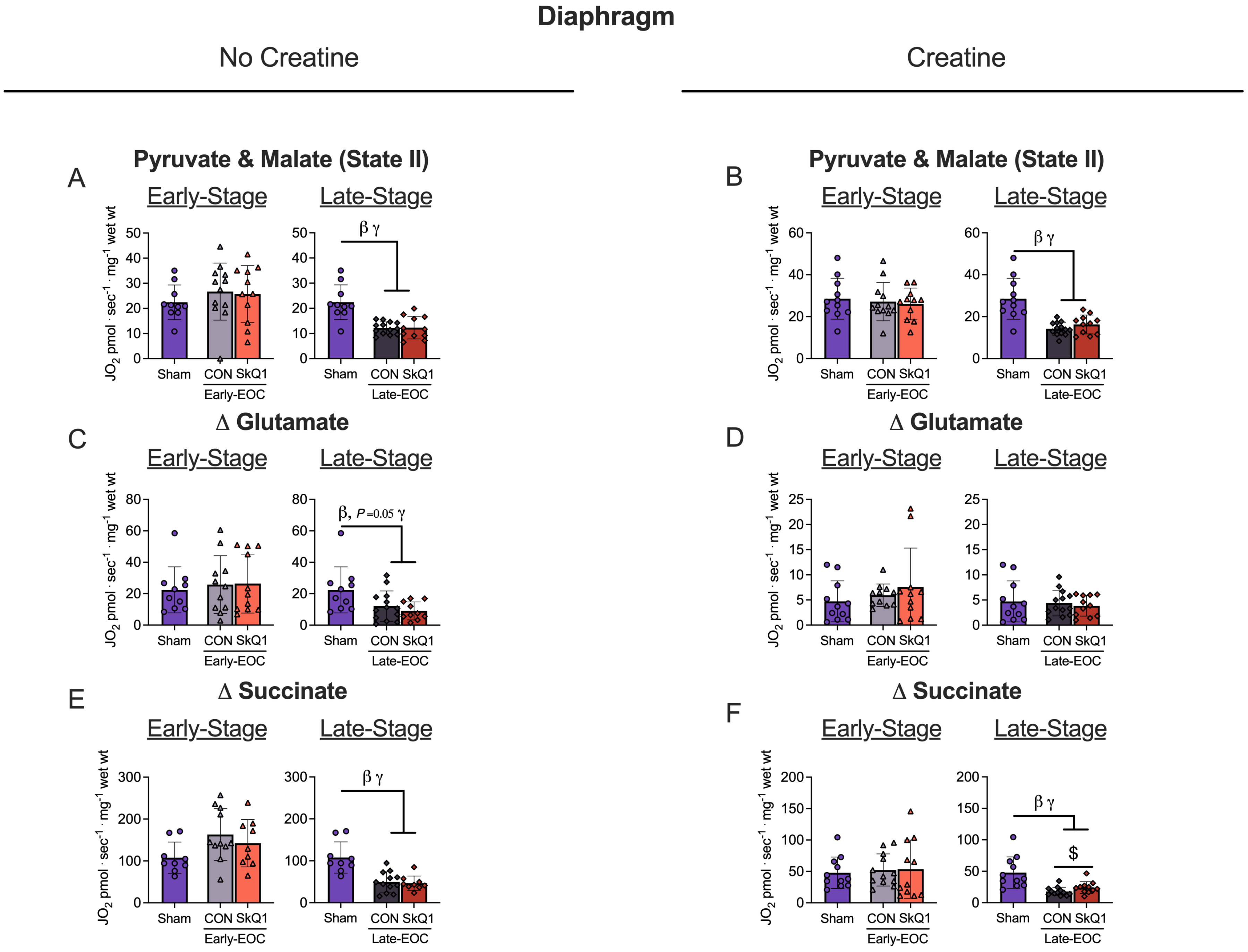


**SFigure 6. Multiple substrate evaluation of oxygen consumption in diaphragm permeabilized muscle fiber bundles.** Oxygen consumption was evaluated in the absence of creatine in Early-Stage and Late-Stage mice when stimulated with pyruvate & malate (**A**), glutamate (**C**) and Succinate (**E**). This was repeated in the presence of creatine (**B, D, E**). All data was analyzed using a one-way ANOVA and followed by a two-stage step-up method of Benjamini, Krieger and Yukutieli multiple comparisons test. Data that was not normally distributed was analyzed with a Kruskal-Wallis test followed by the same post-hoc analysis. Sham Control (Sham), Early-Stage Cancer Control (Early-EOC CON), Early-Stage Cancer SkQ1 (Early-EOC SkQ1), Late-Stage Cancer Control (Late-EOC CON), Late-Stage Cancer SkQ1 (Late-EOC SkQ1). Results represent mean ± SD. n=10-12. ; $\beta$ $P$<0.05 Sham Vs Late-EOC CON; $\gamma$ $P$<0.05 Sham Vs Late-EOC SkQ1.
