## Supplemental Figure 7 for "Mitochondrial-targeted plastoquinone therapy ameliorates early onset muscle weakness that precedes ovarian cancer cachexia in mice"

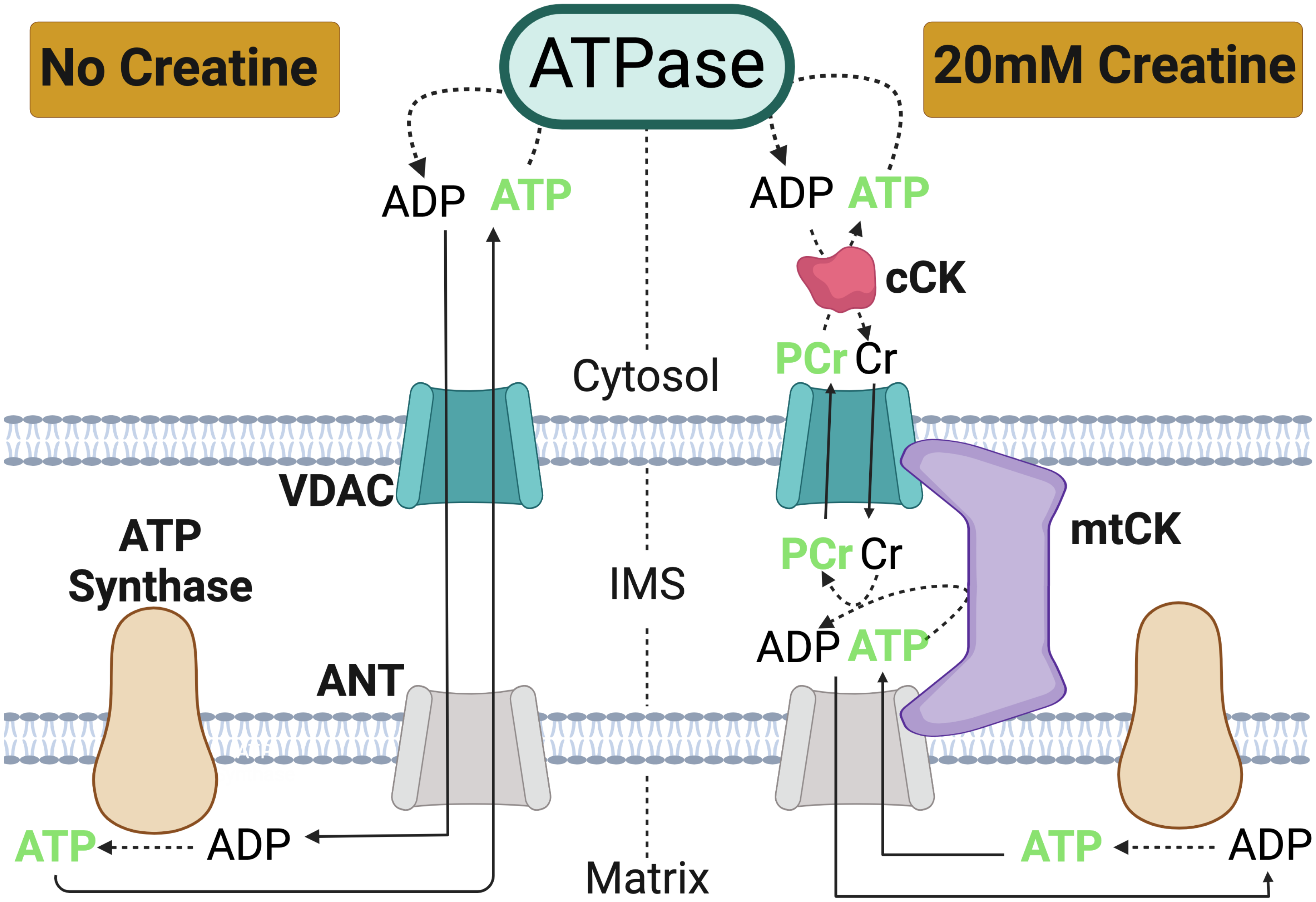


**SFigure 7. Schematic representation of high energy phosphate shuttling in the absence (left) and presence of creatine (right).** ADP generated through various ATPases in the cell is shuttled through the cytosol into the mitochondrial matrix through VDAC and ANT respectively where ADP catalyzes ATP synthase to generate ATP (Left; No Creatine). mtCK accelerates matrix ADP/ATP cycling and ATP synthesis by transferring a high energy-phosphate from ATP to Cr to generate PCR in the IMS. PCr and Cr diffuse rapidly in the cytosol where cCK catalyzes the transfer of high-energy phosphate from PCr to ADP to generate ATP in the cytosol (Right; 20mM Creatine). Voltage dependent anion channel (VDAC); adenine nucleotide translocase (ANT); mitochondrial creatine kinase (mtCK); cytosolic creatine kinase (cCK); phosphocreatine (PCr); Creatine (Cr); Intermembrane space (IMS); adenosine diphosphate (ADP); adenosine triphosphate (ATP). Created with Biorender.
