## Supplemental Figure 8 for "Mitochondrial-targeted plastoquinone therapy ameliorates early onset muscle weakness that precedes ovarian cancer cachexia in mice"

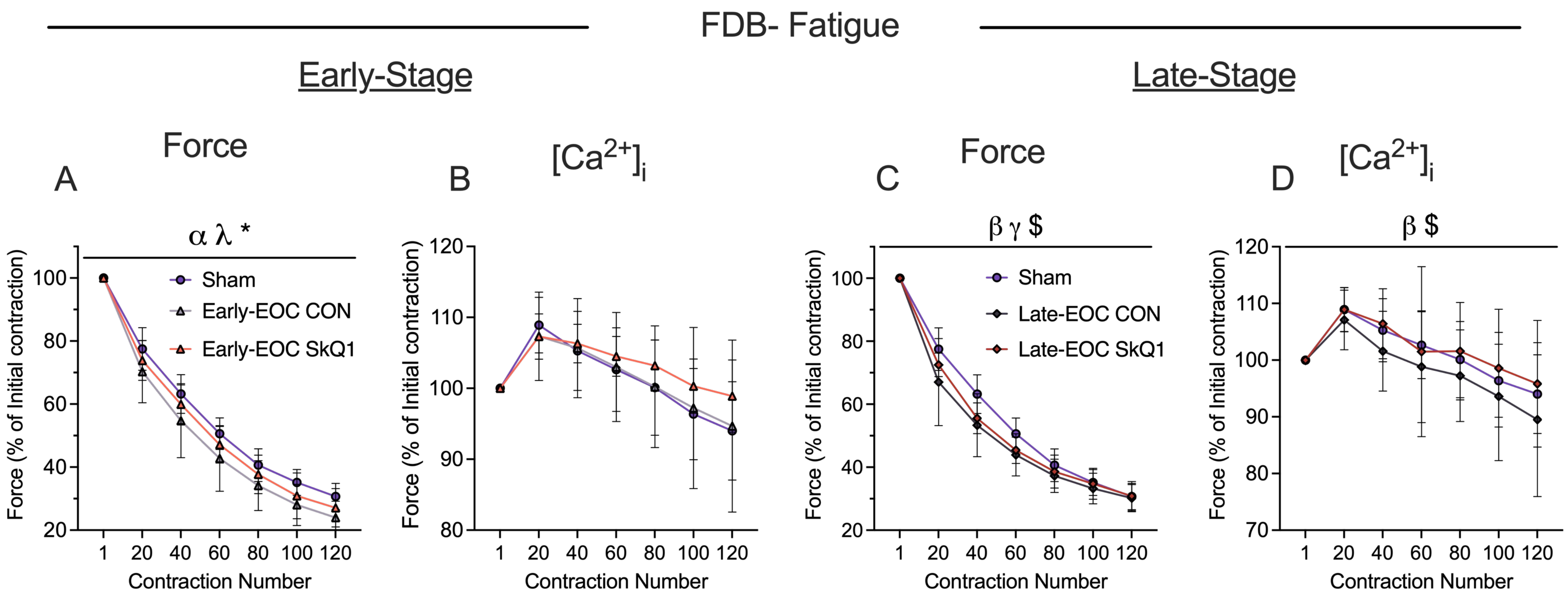


**SFigure 8. Evaluation force-fatigue and calcium-fatigue relationship in FDB single fibers.** In-Vitro FDB fatigue was completed (60Hz for 300ms every second for 120 contractions) to assess the effects of SkQ1 on fatiguability in Early-Stage and Late-Stage mice **(A,C** n=6-10**)**. This was repeated using single fibers for myoplasmic calcium release assessments **(B,D** n=9-12**)**. All data was analyzed using a two-way ANOVA followed by a two-stage step-up method of Benjamini, Krieger and Yukutieli multiple comparisons test. Control Sham Control (Sham), Early-Stage Cancer Control (Early-EOC CON), Early-Stage Cancer SkQ1 (Early-EOC SkQ1), Late-Stage Cancer Control (Late-EOC CON), Late-Stage Cancer SkQ1 (Late-EOC SkQ1), , flexor digitorum brevis (FDB). Results represent mean ± SD. n=10-12. $\alpha$ $P<0.05$ Sham Vs Early-EOC CON; $\lambda$ $P<0.05$ Sham Vs Earl-EOC SkQ1; * $P<0.05$ Early-EOC CON Vs Early-EOC SkQ1; $\beta$ $P<0.05$ Sham Vs Late-EOC CON; $\gamma$ $P<0.05$ Sham Vs Late-EOC SkQ1; $ $P<0.05$ Late-EOC CON Vs Late-EOC SkQ1.
