## Supplemental Figure 9 for "Mitochondrial-targeted plastoquinone therapy ameliorates early onset muscle weakness that precedes ovarian cancer cachexia in mice"

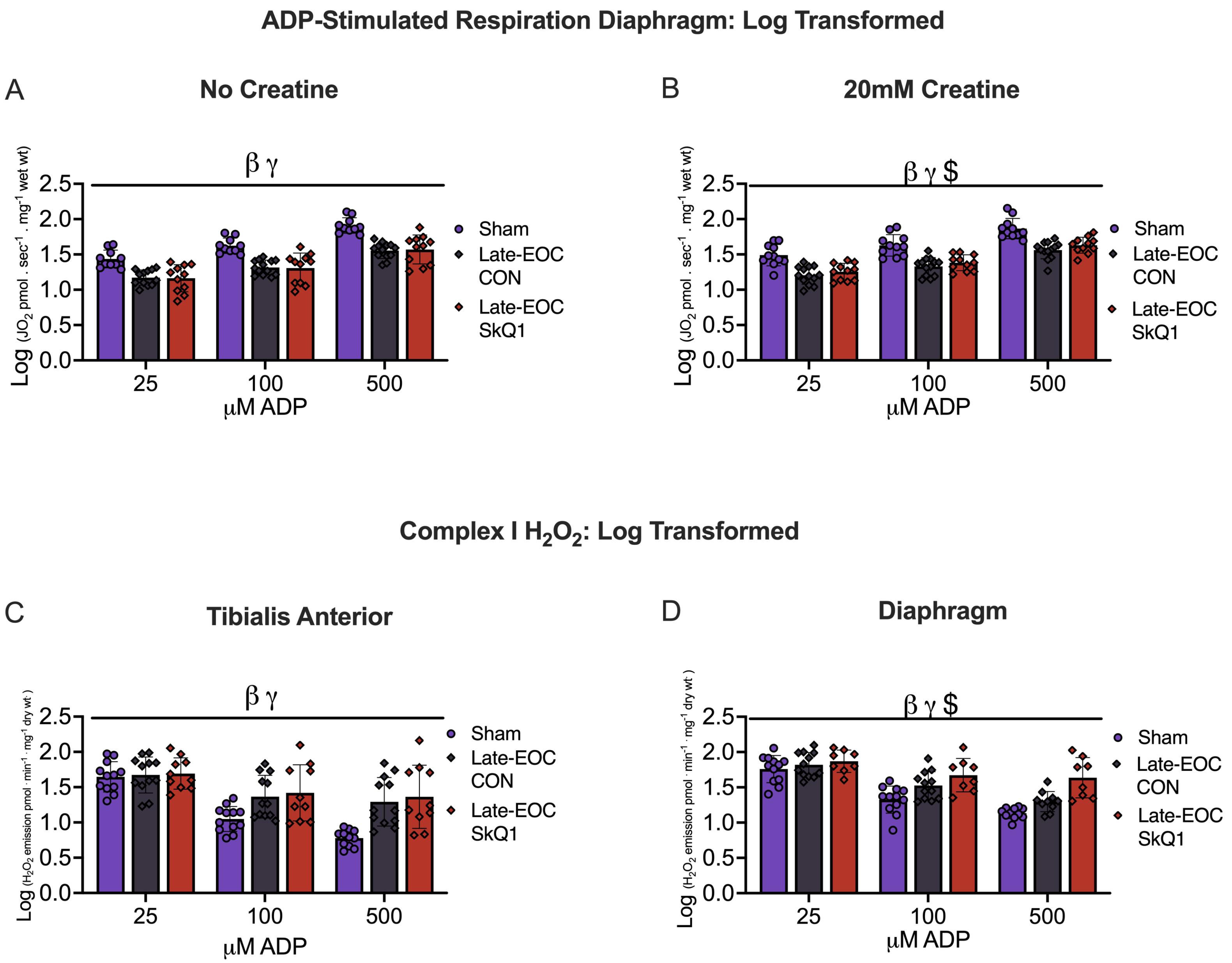


**SFigure 9. Log transformed data.** Data analyzed using a two way ANOVA after log transformation for non-normally distributed data **(A-D)**. Sham Control (Sham), Late-Stage Cancer Control (Late-EOC CON), Late-Stage Cancer SkQ1 (Late-EOC SkQ1), . Results represent mean ± SD. n=10-12. *β* *P*<0.05 Sham Vs Late-EOC CON; *γ* *P*<0.05 Sham Vs Late-EOC SkQ1; $ *P*<0.05 Late-EOC CON Vs Late-EOC SkQ1.
