## Supplemental Table 1 for "Mitochondrial-targeted plastoquinone therapy ameliorates early onset muscle weakness that precedes ovarian cancer cachexia in mice"

### STable 1

| Oligo name | Oligo sequence (5' to 3') |
|---|---|
| m-*actb* Fwd | CATTGCTGACAGGATGCAGAAGG |
| m-*actb* Rev | TGCTGGAAGGTGGACAGTGAGG |
| m-*TNF* Fw | AGAATGAGGCTGGATAAGAT |
| m-*TNF* Rev | GAGGCAACAAGGTAGAGA |
| m-*IL6* Fw | ACAGAAGGAGTGGCTAAG |
| m-*IL6* Rev | AGAGAACAACATAAGTCAGATAC |
| m-*Murf1* Fw | ACCTGCTGGTGGAAAACATC |
| m-*Murf1* Rev | AGGAGCAAGTAGGCACCTCA |
| m-*Atrogin1* Fw | AGCGCTTCTTGGATGAGAAA |
| m-*Atrogin1* Rev | ACGTCGTAGTTCAGGCTGCT |

**STable 1. List of primers used for qtPCR.**
